# Gene regulatory activity associated with PCOS revealed *DENND1A*-dependent testosterone production

**DOI:** 10.1101/2024.05.23.595551

**Authors:** Laavanya Sankaranarayanan, Kelly J Brewer, Graham D Johnson, Alejandro Barrera, Revathy Venukuttan, Ryan Sisk, Andrea Dunaif, Timothy E Reddy

## Abstract

Polycystic ovary syndrome (PCOS) is among the most common disorders affecting up to 15% of the menstruating population globally. It is the leading cause of anovulatory infertility and a major risk factor for type 2 diabetes. Elevated testosterone levels are a core endophenotype. Despite that prevalence, the underlying causes remain unknown. PCOS genome-wide association studies (GWAS) have reproducibly mapped a number of susceptibility loci, including one encompassing a gene regulating androgen biosynthesis, DENND1A. Identifying the causal variants within these loci will provide fundamental insight into the precise biological pathways that are disrupted in PCOS. We report the discovery of gene regulatory mechanisms that help explain genetic association with PCOS in the GATA4, FSHB and DENND1A loci using a combination of high throughput reporter assays, CRISPR-based epigenome editing, and genetic association analysis from PCOS case and control populations. In addition, we found that increased endogenous DENND1A expression causes elevated testosterone levels in an adrenal cell model, specifically by perturbing candidate regulatory elements. These results further highlight the potential for combining genetic variant analyses with experimental approaches to fine map genetic associations with disease risk.

## Introduction

Polycystic Ovary Syndrome (PCOS) is one of the most common disorders affecting people who menstruate with prevalence rates of 5% to 15%^1^, depending on the diagnostic criteria applied^2^. It is the leading cause of anovulatory infertility. PCOS is commonly associated with insulin resistance and obesity, disorders that confer increased risk for type 2 diabetes as well as for other serious cardiometabolic morbidities across the lifespan^3,4^. However, the cause(s) of PCOS remains unknown and the disorder is relatively understudied compared to other common medical conditions affecting women^5^

Genetic factors are a major contributor to PCOS. Twin studies estimate that the narrow-sense heritability of PCOS is ∼79%^6^. There are 30 genomic loci that are associated with increased PCOS risk^7–16^ (GWAS catalog accessed 20 Oct 2023). The associated regions encompass genes involved in neuroendocrine, reproductive, and metabolic pathways. The functional consequences of noncoding genetic variants associated with complex traits such as PCOS have been exceptionally difficult to elucidate^17,18^. One challenge of fine mapping GWAS signals is the difficulty in identifying causal genetic variant(s) from other genetic variants in regions of strong linkage disequilibrium (LD). In general, the lead GWAS SNPs are not the causal variants but are tagging regions of the genome containing non-coding pathogenic variants^17,19^ that contribute to common disease risk by altering regulatory element activity and downstream gene expression^20,21^. Nevertheless, GWAS have provided considerable insight into PCOS causal pathways. DENND1A was first identified as a PCOS candidate gene in GWAS^1^. DENND1A was subsequently shown to be an important regulator of theca cell androgen biosynthesis where ectopic overexpression led to increased androgen production^22–24^. Collectively, rare variants in DENND1A were associated with PCOS quantitative traits in 50% of affected families^24^. Taken together with previous studies indicating that elevated testosterone levels were a consistent endophenotype in sisters of women with PCOS^25^, these genetic analyses implicate DENND1A as a core gene^26^ in PCOS pathogenesis. However, a mechanistic link between the noncoding genome, altered DENND1A expression, and testosterone production has yet to be demonstrated.

The goal of this study was to systematically evaluate the effects of non-coding genomic regions associated with PCOS risk on gene regulatory element activity. To do so, we measured regulatory element activity across PCOS-associated genomic loci and identified genetic variants that alter that activity^27–31^. To measure regulatory activity, we used high-throughput reporter assays because they can quantify the regulatory activity of millions of genomic fragments at once. This scale enables systematic studies of the effects of non-coding variants across megabases of the genome and in many different cell types^27,32–35^. To prioritize variants, we used a combination of allele-specific reporter assays, and targeted genetic association within the identified regulatory elements.

As proof of concept that we the identified regulatory mechanisms contributing to PCOS, we perturbed PCOS-associated regulatory elements near DENND1A using CRISPR-based epigenome editing^36–38^. We found that epigenetic activation of those regulatory elements in an androgen-producing adrenocortical cell model caused both increased DENND1A expression and increased testosterone production. Together, these findings suggest a novel endogenous gene-regulatory mechanism contributing to PCOS; and demonstrate an approach for identifying additional molecular mechanisms of PCOS.

## Results

### Measuring the regulatory activity of PCOS-associated regulatory elements

To identify gene regulatory elements in which genetic variation can contribute to PCOS risk, we analyzed 14 genetic associations identified in cohorts of European and Han Chinese ancestry at the time of this study^9–11,13,14^ (Table 1). Those 14 associations included several genes involved in hormone synthesis via the hypothalamic-pituitary-ovarian axis including FSHR, FSHB, LHCGR and DENND1A. We focused on two human cell models: a testosterone-producing adrenal cell line, H295R; and an estradiol-producing ovarian granulosa cell line, COV434^39–41^.

**Table 1:**
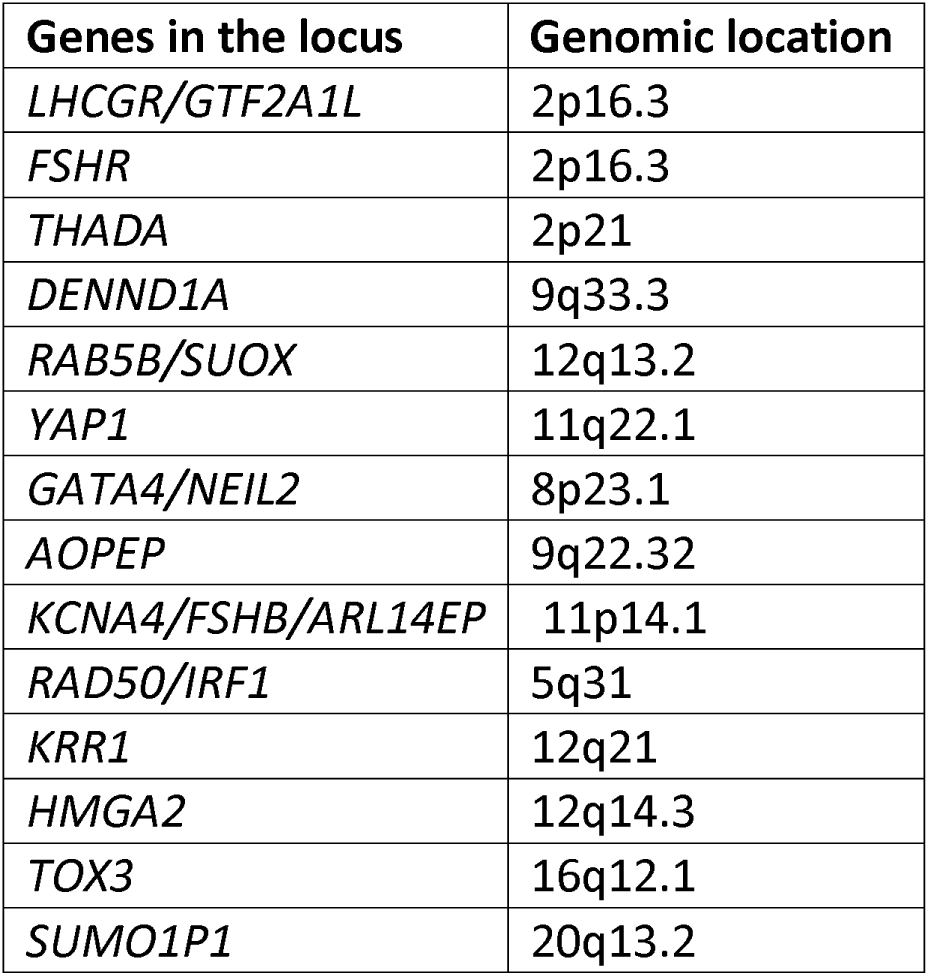
List of PCOS GWAS loci selected for STARR-seq experiments.

To measure regulatory activity in these two cell lines, we used a high-throughput reporter assay known as STARR-seq^29,30^ (Figure 1a). STARR-seq can assay millions of DNA fragments for regulatory activity. STARR-seq assays work through two key libraries – an input library termed ‘assay library’ and a library of the regulatory effect readout termed ‘reporter library’ in this study. Briefly, the assay library consists of plasmid reporter assays containing diverse DNA fragments of interest. When transfected into cells, the DNA fragments regulate their own transcription into mRNA molecules. Thus, by sequencing the reporter library of the resulting mRNA fragments, one can estimate the regulatory activity of each DNA fragment in the assay library.

**Figure 1:**
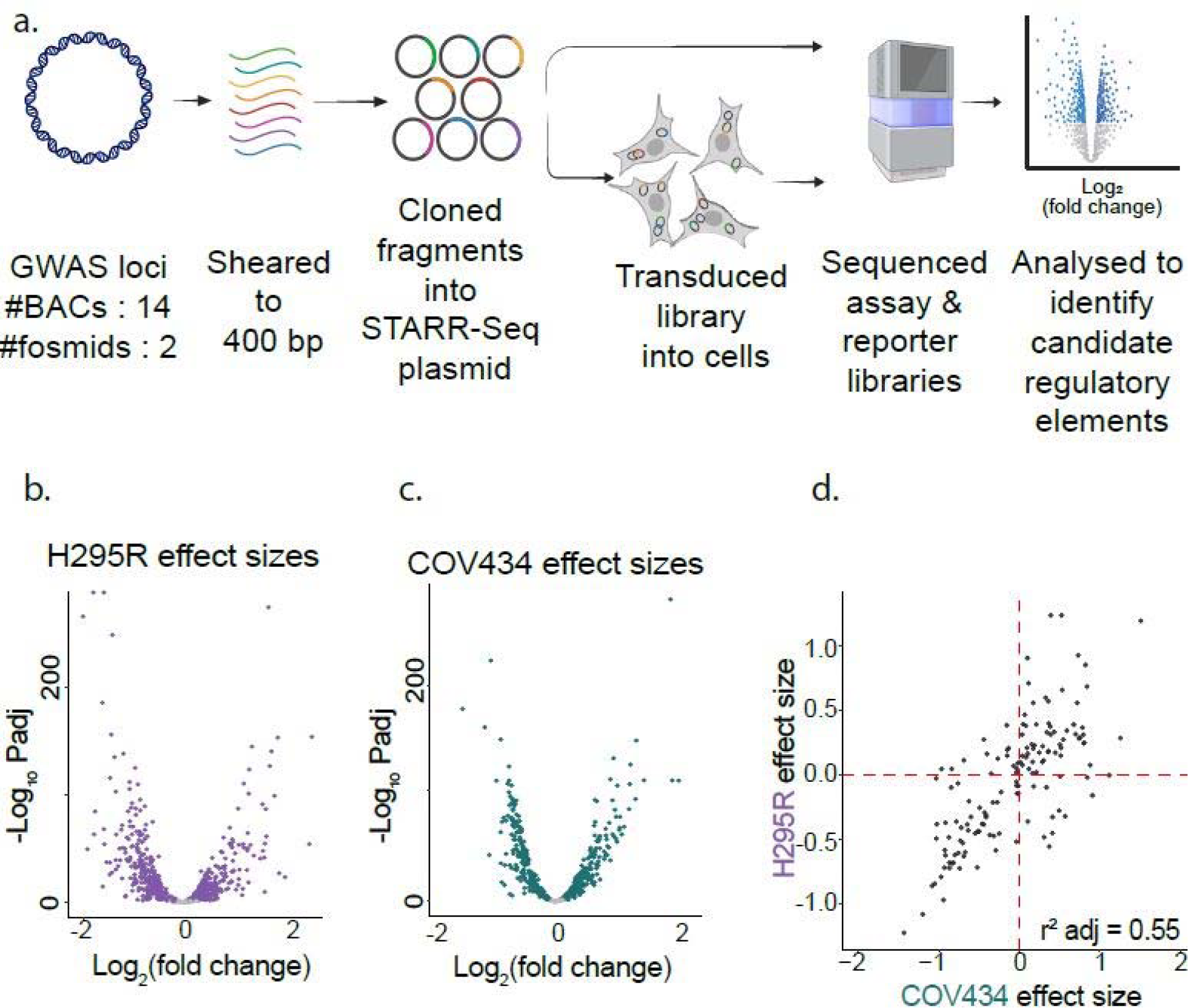
Measuring the regulatory activity in PCOS GWAS loci. a. Overview of targeted STARR-seq method: We selected bacterial artificial chromosomes (BACs) or fosmids spanning 14 PCOS GWAS loci and sheared them to ∼400bp. The sheared fragments were inserted into the digested STARR-seq backbone (Addgene#99296). The resulting plasmid library was sequenced to form the control assay library. For measuring regulatory activity, the plasmid pool was transfected into the respective cell lines (2 μg plasmid pool / 1 million cells). Six hours post transfection, RNA was isolated from the cells, and the STARR-seq transcripts were enriched and sequenced as the output reporter library. Candidate regulatory elements were called using CRADLE^42^ and effect sizes estimated with DESeq2^43^. b. STARR-seq effect size for H295R cells: The effect size is estimated as pseudo log_2_ (fold change) using DESeq2 on CRADLE-corrected STARR-seq peak calls. c. STARR-seq effect size for COV434 cells: The effect size is estimated as pseudo log_2_ (fold change) using DESeq2 on CRADLE-corrected STARR-seq peak calls. d. Comparing effect sizes of shared regulatory elements: About 145 of the regulatory elements were shared between the two cell lines. The adjusted correlation coefficient, r^2^ is 0.55.

We constructed a STARR-seq assay library that spans 14 PCOS GWAS loci and encompassed 2.9 Mb of the human genome (Supplementary Table, S1). The assay library includes 179 open chromatin regions identified in H295R and COV434 (Figure S1). The median fragment length in the assay library was 320 bp, and the 260 bp in the reporter library (Figure S2). Assay library covers the target region at a median of >300x (Figure S3) and replicates are highly correlated with Pearson correlation coefficient (Pearson’ r > 0.95, Figure S4).

We called 956 regulatory elements in the 14 PCOS GWAS loci across the two cell models (Supplementary Tables S2 and S3) at a false discovery rate (FDR) ≤0.5%^42^. Between replicates in the same cell model, the estimated regulatory element activity was highly correlated (0.84 ≤ r ≤ 0.90, Figure S5). Much of the observed variation in effect sizes can be attributed to differences between assay and reporter libraries, and differences between cell lines (Figure S6). The strong correlation suggested that the targeted STARR-seq approach robustly estimated regulatory activity for the cell types within the PCOS GWAS loci.

We identified 464 and 585 regulatory elements in COV434 and H295R cells, respectively. In both cell models, about half of the identified regulatory elements had enhancer activity, and half had repressor activity^43^ (Figure 1b and 1c). There were 93 regulatory elements identified in both cell lines. The regulatory activity of those commonly identified elements was highly concordant. The effect sizes in shared regulatory elements were substantially correlated (Pearson’s r = 0.81, p < 2×10^-16^), and the direction of effects was the same for 85% of shared elements (Figure 1d). The concordance in the direction of effect increased to 93% when we required the regulatory element calls to overlap in the genome by at least 50% (Figure S7, Pearson’s r = 0.85, p < 2×10^-16^). To our knowledge, this data set is the largest reporter-assay screen for enhancers in adrenal and ovarian cell models.

### Regulatory element activity in PCOS GWAS regions corresponds to regions of chromatin accessibility

Enrichment of genetic associations in tissue specific sites of increased chromatin accessibility can predict causal tissues of disease^44,45^. Of the PCOS associated SNPs in the GWAS catalog across the 14 loci we tested, five of those variants overlapped DNaseI hypersensitive sites (DHS) from the ENCODE consortium^46^. To increase confidence that the regulatory elements identified by STARR-seq are active in H295R and COV434 cells, we evaluated whether STARR-seq regulatory elements correspond to chromatin accessibility in the same cell lines. We identified ∼73,000 and ∼66,000 open chromatin sites in H295R and COV434, respectively, using ATAC-seq and MACS2 peak calling with false discovery rate < 0.1. Between 40 and 50% of the open chromatin sites identified in each cell line overlapped sites in the other cell line (Figure S8, S9, S10). Those results revealed that a substantial number of chromatin accessible sites were shared between cell lines.

There were 116 chromatin accessible sites within the 14 genomic regions we assayed with STARR-seq across each COV434 and H295R cell lines. Of those, 39 (34%) and 37 (32%) had regulatory activity in H295R and COV434 cells, respectively, according to STARR-seq assays (Supplementary Tables S4 and S5). For H295R cells, the overlap between chromatin accessibility and STARR-seq activity was ∼4-fold more than what would be expected if STARR-seq sites were randomly distributed across the genomic regions. For COV434, the overlap was ∼6-fold more than expected by random (Fisher’s exact test, p < 2 x 10^-4^ for each). There was also significantly greater regulatory activity in open chromatin regions in the same cell type or tissue than in regions with less chromatin accessibility (Figure 2a) (Mann-Whitney U, p < 10^−10^ for H295R, p <0.01 for COV434). Conversely, there was more chromatin accessibility in regions where we identified regulatory element activity (Figure 2b).

**Figure 2:**
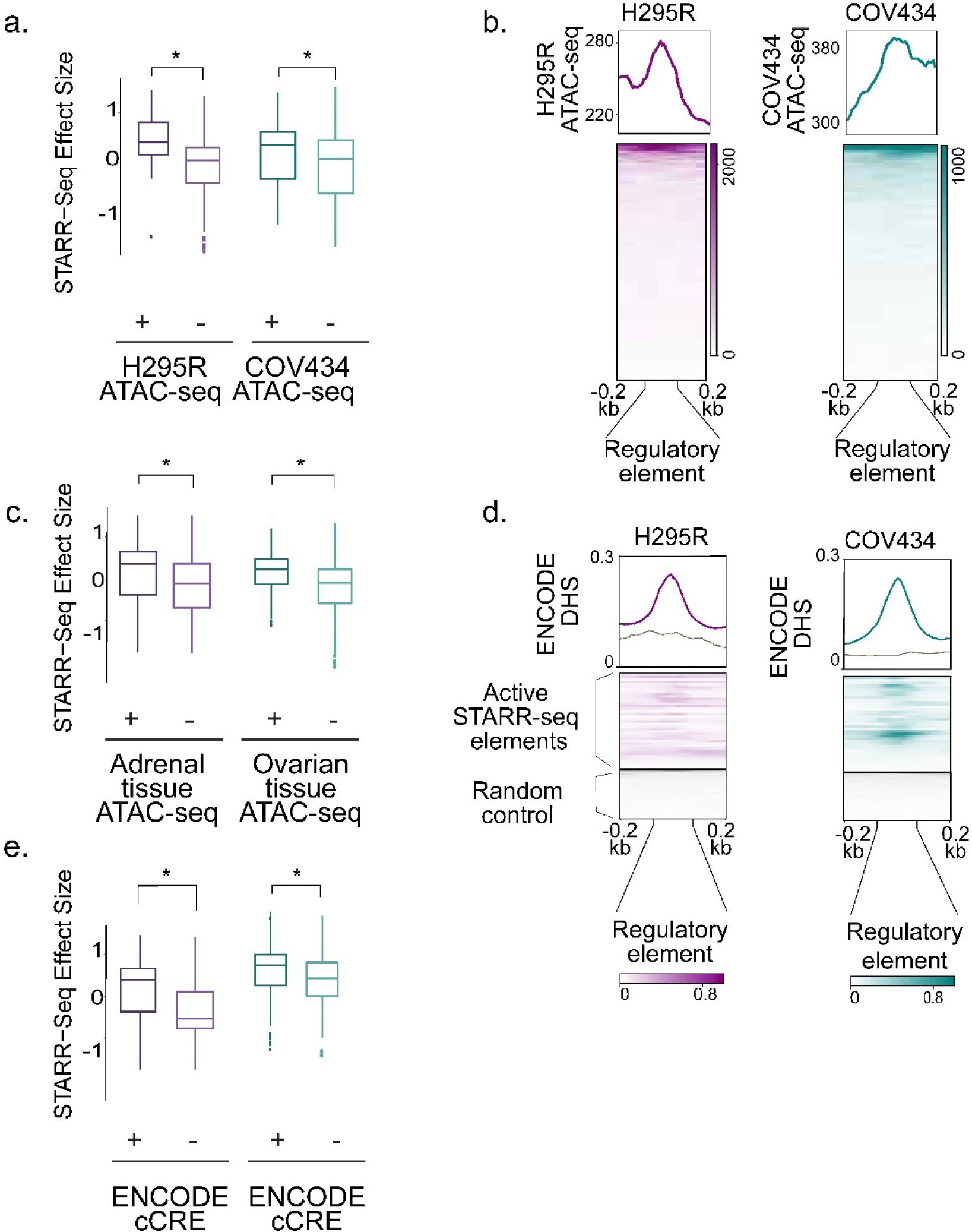
Characterizing candidate regulatory elements. a,c,e. Candidate regulatory elements in H295R cells and COV434 cells with increasing activity correspond to regions with increased evidence of functionality. STARR-seq regulatory activity is measured across overlap with the respective cell line ATAC-seq (a), GTEx primary tissue ATAC-seq (c) and ENCODE candidate cis-regulatory elements (e). Candidate regulatory elements identified in H295R (purple track) and COV434 (teal track) cell lines. Each STARR-seq track is reported as assay (input) subtracted reporter (output) libraries. Boxes correspond to regulatory elements overlapping accessible chromatin regions (grey). b. Aggregate profile plots of chromatin accessibility based on ATAC-seq on the respective cell lines centred on the candidate regulatory elements (with increasing and decreasing effect sizes) across 400 bp windows for both cell lines (H295R in purple, COV434 in teal). d. Aggregate profile plots of chromatin accessibility based on ENCODE DNaseI Hypersensitive sites (DHS) centred on the active candidate regulatory elements across 400 bp windows for both cell lines (H295R in purple, COV434 in teal). Control regions (grey) are randomly generated genomic regions that are chromosome-, length- and GC-matched to the STARR-seq elements.

We also investigated similarities and differences in regulatory activity between H295R and COV434 cells. There were 69 genomic regions that had significant regulatory activity and significant chromatin accessibility in either cell model. Of these, seven had regulatory activity in both cell models. The small overlap was due to differences in statistical power. Specifically, regulatory activity was similar across both cell types (ρ = 0.65, Figure S11). There was also no strong evidence of elements with opposing regulatory activity between cell types. Taken together, the high concordance of regulatory effect size across STARR-seq in H295R and COV434 suggested that regulatory activity was largely similar between the two steroidogenic cell lines.

To relate cell line observations to the corresponding primary tissues, we evaluated if STARR-seq regulatory activity was enriched in chromatin accessible sites in adrenal and ovarian tissues^46,47^. Approximately 18% of the identified H295R regulatory elements overlapped with open chromatin from primary adrenal tissue, and 24% of the identified COV434 regulatory elements overlapped with open chromatin from primary ovarian tissue. The overlap was a 2.8 and 3.1-fold enrichment in H295R and COV434, respectively, over what would be expected if regulatory elements were randomly distributed across the assayed regions (Fisher’s exact test, p-value < 10^−7^ for both). As with our observations in H295R and COV434 cells, regulatory activity was greater in regions of accessible chromatin in primary tissue compared to those without accessible chromatin (Figure 2c, Mann-Whitney U, p < 10^−9^). This result indicated that regulatory activity measurements in H295R and COV434 cells corresponded to activity in primary adrenal and ovarian cells, respectively.

Regulatory activity in H295R and COV434 cells also corresponded to chromatin accessibility in other tissues. About 50% of the regulatory elements we identified via STARR-seq (n = 296 for H295R, n = 304 for COV434) overlapped chromatin accessible sites identified in diverse tissues as part of the ENCODE project^46,47^. The overlap was 1.7- and 2.7-fold enriched over what would be expected if regulatory activity was randomly distributed across the assayed regions in H295R and COV434, respectively (Figure S10, Mann-Whitney U, corrected p-value < 10^−4^). ENCODE DNase hypersensitive sites also had increased activity in STARR-seq regulatory elements (Fisher’s exact test p < 10^−12^, Figure 2d, Figure S11). We observed similar results when focusing on enhancer-like regions defined across diverse cells and tissues by the ENCODE project^48^. Specifically, ∼30% of the regulatory elements we identified overlaped proximal or distal enhancers defined by ENCODE (n = 158 for H295R; n = 207 for COV434); and quantitative estimates of regulatory activity was greater in regions identified as enhancer-like sequences (Figure 2e).

### PCOS-associated genetic variants fine-mapped to within regulatory elements

To discover genetic variants that may alter regulatory activity and gene expression, we completed genetic association analyses focused on the regulatory elements we identified (Figure 3a). To identify additional risk variants within these functional regulatory elements, we first tested for genetic associations between single nucleotide polymorphisms (SNPs) with minor allele frequency (MAF) >1% and PCOS disease within the regulatory elements we identified. Across a cohort of 983 PCOS cases and 2951 controls^9^, we tested 759 SNPs in H295R cells, and 486 in COV434 cells. We found 19 variants that were significantly associated with PCOS at an adjusted p-value (Bonferroni correction) threshold of 1.15 x 10^-4^ – 2.55 x 10^-4^ (Table 2). Of the associated variants, four were in the follicle stimulating hormone subunit beta (FSHB) locus, six were in the neighboring ARL14EP-DR locus and two were in the GATA4/NEIL2 locus (Figure 3b and 3c, Table 2, Supplementary workbook). There were four previously identified PCOS-associated risk variants in the regulatory elements we assayed: rs6022786 is an intergenic variant near SUMO1P1; rs2268361 is a variant in an intron of FSHR; rs11225154 is a variant in an intron of YAP1 and rs10835638 is a variant in an intron of ARL14EP-DT ^8,11,13^. Of those, only rs6022786 was tested in this analysis, and there was not a significant association with PCOS in our cohort.

**Figure 3:**
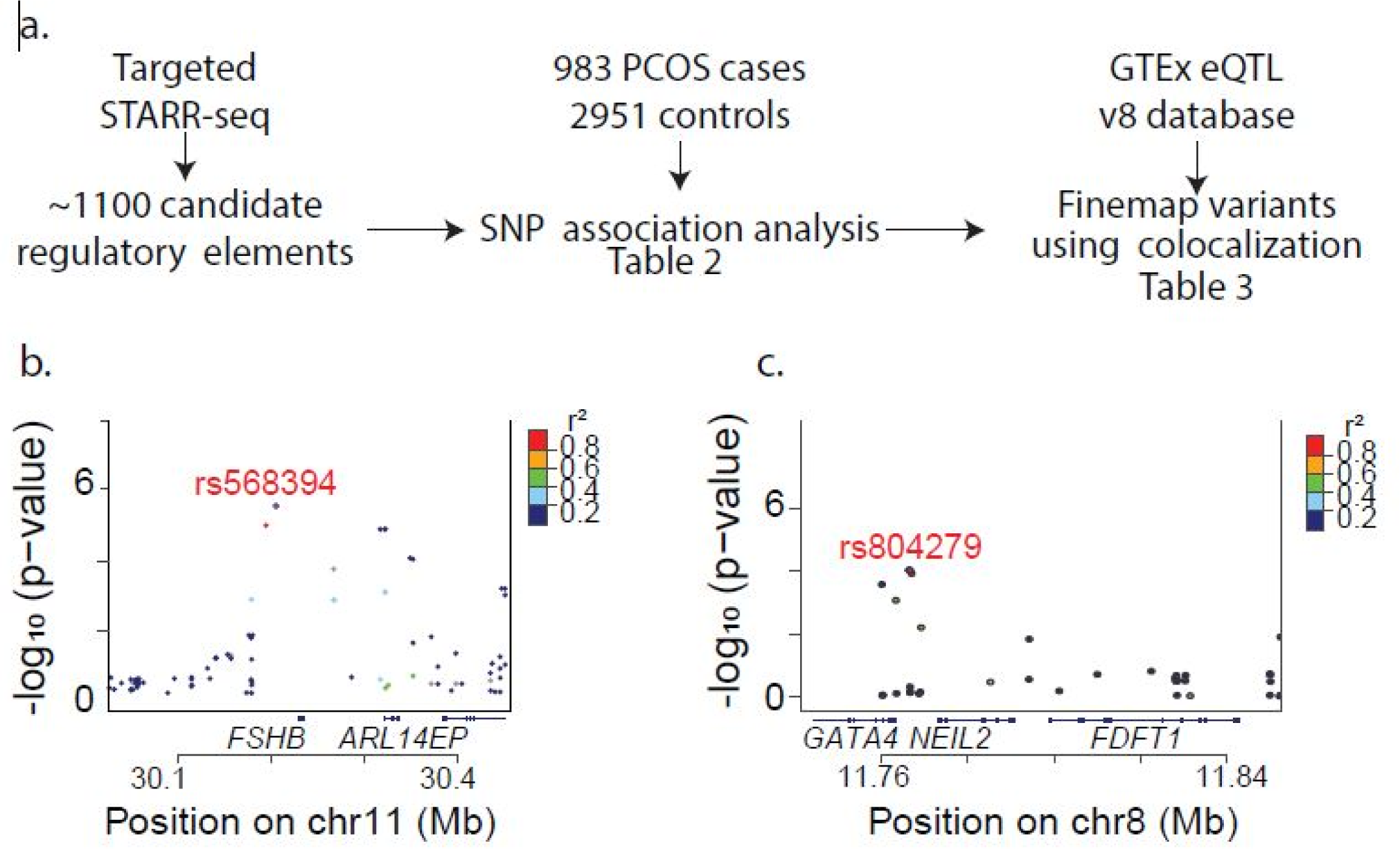
Prioritizing PCOS-associated variants within functional regulatory elements. a. Association analysis to identify PCOS-associated variants in regulatory elements. We use the candidate regulatory elements from STARR-seq experiments to define the genomic regions of interest. We then performed an association analysis to identify variants associated with PCOS using a cohort of 983 PCOS cases and 2951 controls (results in Table 2). We then colocalized the association analysis results with GTEx eQTL SNPs to identify SNPs and genes as those likely involved in PCOS pathogenesis (results in Table 3). b,c. Locuszoom plots for PCOS-associated variants in candidate regulatory elements in FSHB/ARL14EP locus (b) and GATA4/NEIL2 locus (c).

**Table 2:** Top variants associated with PCOS within STARR-seq regulatory elements. OR = odds ratio, MAF = minor allele frequency.

| CHR | BP | OR | P | MAF | rsID |
| --- | --- | --- | --- | --- | --- |
| 11 | 30205987 | 1.452 | 3.08E-06 | 0.4044 | rs568394 |
| 11 | 30195456 | 1.419 | 1.16E-05 | 0.4005 | rs574096 |
| 11 | 30317914 | 1.544 | 1.49E-05 | 0.1676 | rs7929660 |
| 11 | 30323044 | 1.544 | 1.49E-05 | 0.1676 | rs4071558 |
| 11 | 30323178 | 1.544 | 1.49E-05 | 0.1676 | rs4071559 |
| 11 | 30350068 | 1.485 | 1.05E-04 | 0.1605 | rs12363432 |
| 11 | 30353061 | 1.465 | 1.14E-04 | 0.1754 | rs7949790 |
| 11 | 30213950 | 1.452 | 2.58E-06 | 0.4189 | rs586326 |
| 11 | 30295275 | 1.545 | 1.43E-05 | 0.1675 | rs12278989 |
| 11 | 30347873 | 1.376 | 5.37E-05 | 0.4513 | rs6484481 |
| 11 | 30359529 | 1.472 | 8.99E-05 | 0.1766 | rs10835661 |
| 8 | 11766380 | 1.408 | 9.20E-05 | 0.2597 | - |
| 11 | 30233136 | 1.359 | 9.77E-05 | 0.4534 | rs594982 |
| 11 | 30227279 | 1.358 | 1.00E-04 | 0.4517 | rs1782509 |
| 11 | 30227537 | 1.357 | 1.04E-04 | 0.4532 | rs1716024 |
| 11 | 30237442 | 1.357 | 1.07E-04 | 0.4527 | rs615577 |
| 11 | 30199192 | 0.7338 | 1.15E-04 | 0.3929 | - |
| 8 | 11766907 | 1.405 | 1.18E-04 | 0.2518 | rs34928882 |
| 11 | 30341554 | 1.488 | 9.67E-05 | 0.1601 | rs7926666 |

To relate these variants to effects on gene expression, we next tested for colocalization^49^ between PCOS-associated genetic variation in regulatory elements and expression quantitative trait loci (eQTLs) from GTEx^50^. Specifically, we used significant single tissue-eQTL association for this analysis. We identified seven variants in seven different loci where PCOS association and gene expression association colocalized with a posterior probability > 0.6 (Table 3, Supplementary Table S11). We further refined this analysis to only include adrenal and ovarian eQTLs. Focusing on these tissues has the advantage of being relevant to specific PCOS mechanisms but also reduced statistical power because there were fewer samples. Using this tissue-specific analysis, we identified four of the same variants (Supplementary Table S11). Two of the variants – rs804271 in the GATA4/NEIL2 locus, and rs11349741 in the FSHB/ARL14EP locus – were also significant in the genetic association focused on regulatory elements described above.

**Table 3:** Colocalization of PCOS-associated variants with eQTL data from GTEx. We identified 7 variants with high probability of likely causal of both PCOS-risk-association and altered gene expression from eQTL data. Colocalization was done using coloc (R package). PP.AllTissue is the probability of colocalization using eQTLs from all tissues in GTEx.

| Variant position | PP.AllTissue | Gene(s) associated from eQTLs |
| --- | --- | --- |
| chr8:11769705 | 1 | *GATA4* |
| chr12:55999553 | 1 | *RPS26/RAB5B/SUOX* |
| chr20:53949370 | 0.86 | *BCAS1* |
| chr9:123707451 | 0.84 | *DENND1A* |
| chr2:48730442 | 0.76 | *LHCGR* |
| chr11:30341402 | 0.71 | *ARL14EP/FSHB* |
| chr9:94907391 | 0.69 | *C9orf3/AOPEP* |

There are plausible biological mechanisms by which the expression of genes in each locus might affect PCOS pathogenesis. GATA4 codes for a transcriptional factor that is involved in embryogenesis. Functional studies showed that deletion of GATA4 resulted in decreased fertility and granulosa and theca cell proliferation^51^. FSHB is a key peptide hormone that regulates follicular development and is, thus, a strong candidate gene for PCOS. Together, these analyses further fine map genetic variants that alter regulatory activity and the expression of several genes implicated in the development of PCOS.

### Active STARR-seq regions have increased conservation score

Evolutionary conservation is another indicator of biological function that is complementary to chromatin accessibility and STARR-seq analyses. We anticipated that genes affecting fertility would have strong evolutionary consequences. In support of this notion, previous studies have reported that conservation of regulatory elements corresponds to a greater functional role in the organism^52^. Therefore, we investigated patterns of conservation across the regulatory elements we identified. We compared conservation scores of regulatory elements that we identified by STARR-seq across 20 vertebrate species^53^. The STARR-seq regulatory elements with enhancer activity had increased conservation score when compared to GC- and length-matched regions on the same chromosome (Figure S13b, Mann-Whitney U, p < 0.001). We also observed that the accessible chromatin region identified by ATAC-Seq within COV434 and H295R cells have higher conservation scores (Figure S14) when compared to similarly matched genomic regions from the same chromosome. These results further corroborated the functional importance of the regulatory elements we identified.

### Allele-specific regulatory variants identified in DENND1A locus

As proof of concept that the regulatory elements we identified were relevant to PCOS pathogenesis, we focused on mechanisms contributing to altered expression of genes in the DENND1A locus. DENND1A is a guanine nucleotide exchange factor involved in clathrin-mediated endocytosis^54,55^. DENND1A expression has been implicated in androgen biosynthesis ^22–24^, including in H295R cell model^56^. Therefore, we focused on understanding the role of regulatory elements in controlling DENND1A expression and activity in H295R cells.

The DENND1A locus has been reproducibly associated with PCOS in Han Chinese and European cohorts^9,10,24,57–59^. However, the functional variants within DENND1A have not been identified. We mapped 38 candidate regulatory elements between the second and sixth introns of DENND1A spanning ∼180 kb of the genome. Several of these regulatory elements overlapped regions called as candidate cis regulatory elements (cCRE) through ENCODE, or were in regions with increased chromatin accessibility in H295R and COV434 (Figure 4a). The lead GWAS risk variants did not overlap the regulatory elements we identified in this study. However, for most of these candidate regulatory elements, there were common variants in linkage disequilibrium (Figure S15) with the lead GWAS SNPs. Taken together, these results suggested that regulatory variants within candidate regulatory elements could contribute to PCOS pathogenesis by affecting gene expression of the target gene of that regulatory element.

**Figure 4:**
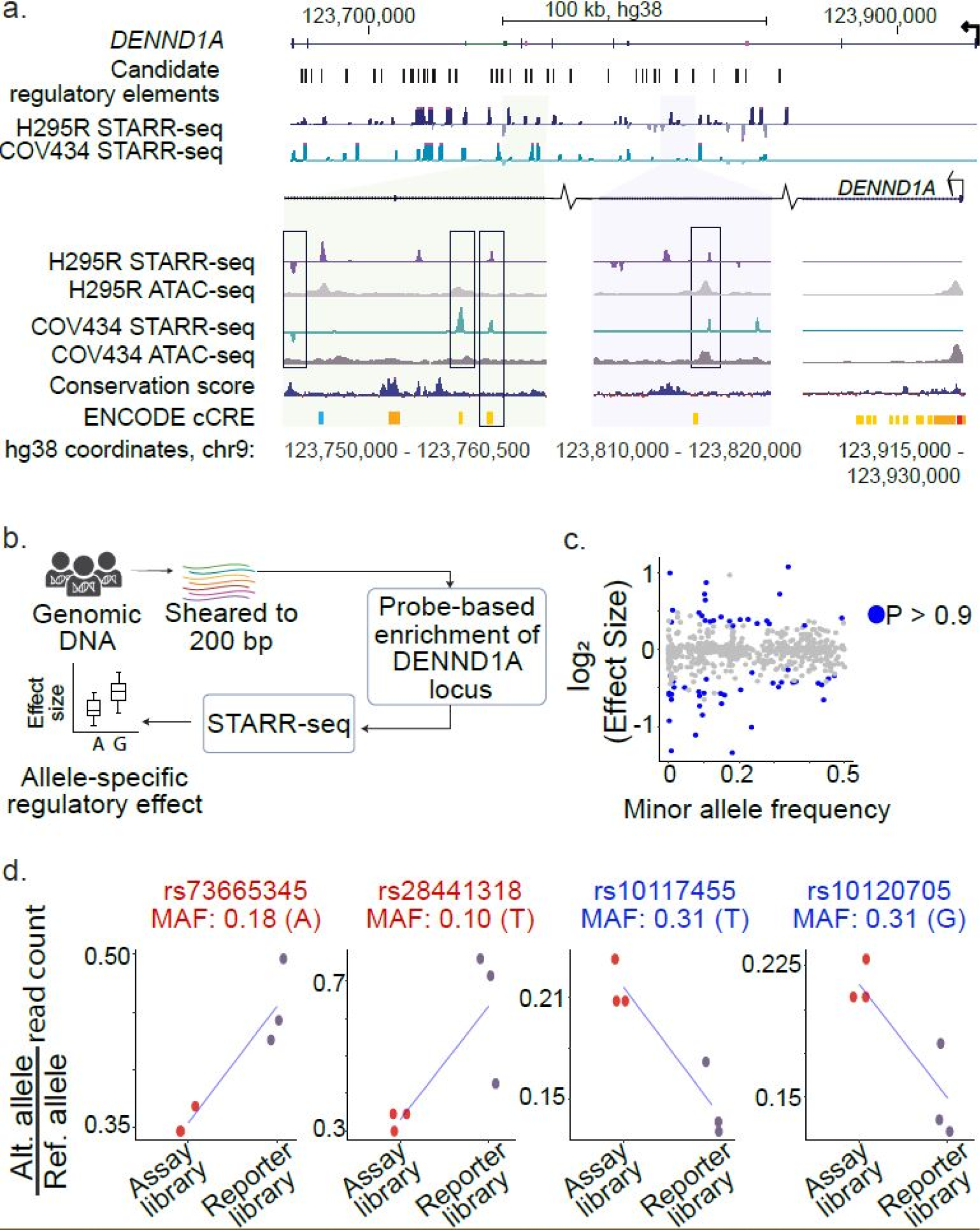
Fine-mapping variants identified four regulatory variants that are also eQTLs for DENND1A. a. Candidate regulatory elements in DENND1A locus identified in H295R (purple track) and COV434 (teal track) cell lines. Each STARR-seq track is reported as assay (input) subtracted reporter (output) libraries. Zoomed-in genomic regions show the candidate regulatory elements, ATAC-seq data conservation score and ENCODE cCREs of that region in detail. b. Overview of enriched DENND1A-STARR-seq method. Genomes from five individuals from the 1000 Genomes Project were sheared to 200 bp. We enriched the target DENND1A locus using RNA-probes (using Agilent Sure Select System). The custom probes were designed to span the DENND1A locus at 2x tiling density. These enriched fragments were then subject to the STARR-seq protocol mentioned in Figure 1a. Allele-specific regulatory effect was estimated using the BIRD model. c. Distribution of allele-specific effect sizes as estimated by BIRD against their minor allele frequencies. The estimated significant SNPs with allele-specific regulatory activity (P > 0.9) are in blue. d. Four variants identified to have allele-specific regulatory activity in H295R. SNPs in red are examples where the alternate allele has increased regulatory activity while SNPs in blue are examples where the reference allele has increased regulatory activity.

To measure the effects of genetic variation across DENND1A on gene expression, we captured genomic DNA spanning the entire DENND1A gene region from three individuals of European ancestry and two of Han Chinese ancestry. We then measured allele-specific regulatory activity in H295R cells using STARR-seq^33^ (Figure 4b, Supplementary Table, S6). In total, we assayed ∼700,000 unique ∼160 bp DNA fragments (Figure S16, Figure S17). The assay library covered the DENND1A gene locus at a median coverage of 140x. The assay libraries were highly concordant, while the measure of log fold change between the reporter and assay libraries was moderately concordant among replicates (Pearson’s r > 0.72, Figure S18).

To estimate the allele specific regulatory effects, we used a Bayesian approach, BIRD, that identifies differences in the relative abundance of alleles in the assay library and in the expressed reporter library^60^. Of the 623 variants we assayed in the targeted locus, 62 had allele specific regulatory activity with a posterior probability, Preg > 0.90 (Supplementary Table S7). On average, the identified variants altered regulatory activity by 40% (Figure S19), and the minor alleles more often had less regulatory activity (chi-squared = 6.9, p-value = 0.009).). We observed a modest correlation between the absolute effect size and the minor allele-frequency of the selected variants as determined by the 1000 Genomes project (ρ = -0.36, p = 0.005, Figure 4c)^61^.

Of the 62 identified regulatory variants we identified, 24 were eQTLs for DENND1A (n = 11) or flanking genes CRB2, RABGAP1 or STRBP (n = 14)^50^. Of those variants, 12 also overlapped open chromatin sites or candidate cis-regulatory elements identified by ENCODE (Figure 4d, Figure S20, Table 4, Supplementary Table S8). Furthermore, the lead variant from colocalization analyses, rs10117940 (Table 5) was also identified in allele-specific analysis with an effect size of 1.299 (p = 0.731). The variant, rs10117940, was in LD with two STARR-seq regulatory variants (rs28441318 and rs73665345) and a PCOS-associated rare variant^24^ (rs78012023) (0.32 < r^2^ < 0.65; 0.5 D’ > 0.9). These findings suggested that several loci within the DENND1A gene contributed to PCOS phenotypes by altering DENND1A gene expression.

**Table 4:** Candidate allele-specific regulatory variants identified. The effect size is measured using the BIRD model (see methods). MAF = minor allele frequency. Posterior probability is the probability of that variant being a regulatory element. Overlap with orthogonal genomic data indicates whether that variant is present in open chromatin as measured by ATAC-seq or DNase-seq, in proximal enhancer like elements (pELS) or distal enhancer like elements (dELS) from ENCODE. The eQTL gene are the gene(s) that variant is associated with as an eQTL from GTEx data.

| Variant position | rsID | Effect size | MAF | Posterior probability | Overlap with orthogonal genomic data | eQTL gene |
| --- | --- | --- | --- | --- | --- | --- |
| chr9:123399942 | rs4466467 | 0.56 | 0.096 | 0.97 | DNase-seq | *CRB2* |
| chr9:123356770 | rs10985986 | 1.32 | 0.226 | 0.96 | pELS | *CRB2* |
| chr9:123930320 | rs10117455 | 0.71 | 0.312 | 0.99 | ATAC-seq | *DENND1A* |
| chr9:123930321 | rs10120705 | 0.73 | 0.311 | 0.98 | ATAC-seq | *DENND1A* |
| chr9:123922454 | rs28441318 | 1.66 | 0.102 | 0.96 | ATAC-seq | *DENND1A* |
| chr9:123708987 | rs73665345 | 1.24 | 0.183 | 0.95 | dELS | *DENND1A* |
| chr9:123679118 | rs10760295 | 0.77 | 0.454 | 0.91 | dELS | *DENND1A* |
| chr9:123281731 | rs10760271 | 0.81 | 0.237 | 0.91 | dELS | *RABGAP1* |
| chr9:123338729 | rs10114139 | 0.73 | 0.435 | 0.91 | dELS | *STRBP* |
| chr9:123358280 | rs62579936 | 1.30 | 0.104 | 0.91 | dELS | *STRBP, CRB2* |
| chr9:123359685 | rs10156609 | 1.57 | 0.104 | 0.99 | pELS | *STRBP, CRB2* |
| chr9:123391031 | rs2491351 | 0.79 | 0.471 | 0.97 | dELS | *STRBP, CRB2, DENND1A* |

### Activation of PCOS-associated regulatory elements increased DENND1A expression

Estimating the effect of regulatory elements on altering gene expression can provide an insight into the underlying mechanisms that contribute to the development of PCOS. While reporter assays like STARR-seq can functionally test for allele-specific regulatory activity, the approach does not identify the target genes of those regulatory elements because the plasmids are not integrated in the genome. One approach to identify target genes of candidate regulatory elements is by epigenomic perturbation of that element. Specifically, a fusion of catalytically inactive Cas9 (dCas9) and histone acetyltransferase domain of P300 is targeted to candidate regulatory elements to measure the effects on the expression of nearby genes^35^. Several studies have demonstrated that dCas9-P300 can act over tens of kilobases, thus allowing the identification of distal gene regulatory elements^37,62,63^.

To identify target genes of PCOS-associated gene regulatory regions, we created dCas9-P300-expressing H295R cells. We targeted dCas9-P300 to four candidate regulatory elements within the introns of the DENND1A gene and to the DENND1A promoter labeled “element 1-4” (Figures 5a, S21). We prioritized those four regulatory elements based on their proximity to DENND1A gene, the strength of STARR-seq signal, chromatin accessibility, and the ability to design guide RNAs (gRNAs) targeting dCas9-P300 to the element (Figure 5a, Supplementary Table, S9).

**Figure 5:**
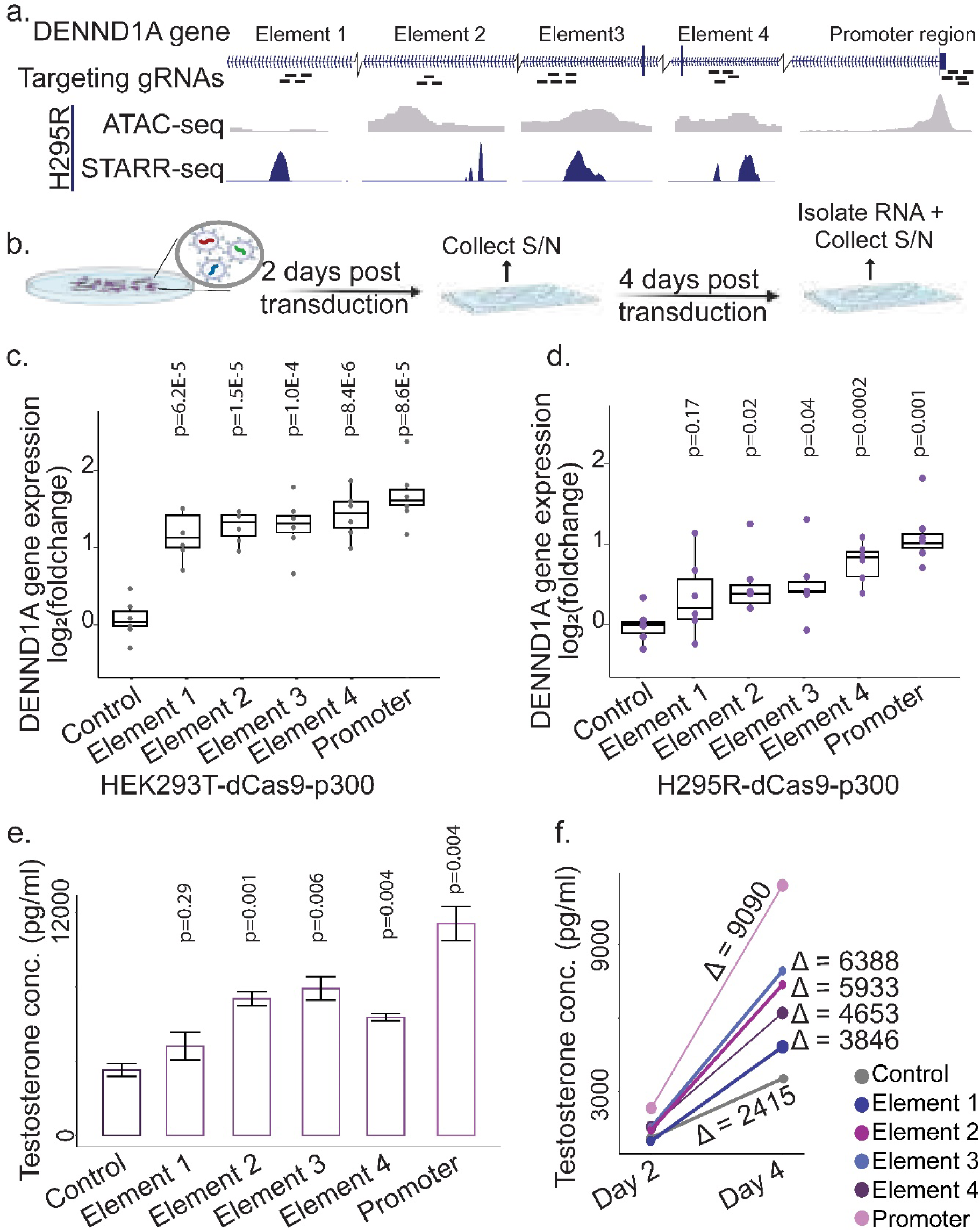
Perturbation of regulatory elements in DENND1A impacts testosterone levels. a. Perturbation loci: We targeted four candidate regulatory elements (Elements 1-4, also Figure S17) and the promoter regions of DENND1A. We designed between 5&7 guides per region. The STARR-seq activity (purple) and chromatin accessibility (grey) for these regions are shown. b. H295R cells that stably express dCas9-p300 were transduced with lentiviral pools of guide RNAs for each regulatory element. The cell media supernatant was collected 2- and 4-days post transduction for measuring testosterone concentration produced by the cells. RNA was harvested from the cells 4-days post transduction to measure changes in gene expression. c, d. Log-fold change of DENND1A expression (GAPDH as control) for HEK293T (c.) and H295R (d.) cells stably expressing dCas9-p300. A set of 5 non-targeting control guides were designed to not target any part of the human genome as control cell population. e. Testosterone concentration (pg/ml) measured in the cell media 4 days post transduction in H295R-dCas9-p300 cells targeted with the specific guide RNA. f. Change in testosterone concentration (pg/ml) between day 2 and day 4 post transduction for each H295R-dCas9-p300 cells targeted with the specific guide RNA. The change in testosterone concentration between the two measurements is represented as Δ (in pg/ml).

We designed 5-7 guide RNAs (gRNAs) for each of the four regulatory elements and promoter region (Supplementary Table, S10). As a negative control, we also designed five guide RNAs to not target any location in the human genome. We made lentiviral pools for each of the four targeted regions and for the negative controls. We then transduced each lentiviral pool into two cell lines, H295R and HEK293T, that were modified to express dCas9-P300. DENND1A was already expressed in both cell lines (average TPM: 20.4 for HEK23T and 15.4 for H295R) ^64,65^, indicating that the gene was not in heterochromatin and thus could be targeted by dCas9-P300 effectively. Finally, we measured the effects on DENND1A expression via qPCR, and levels of testosterone at two time points (Figure 5b).

In HEK293T-dCas9-P300 cells^37^, targeting dCas9-P300 to the DENND1A promoter increased DENND1A expression by 3.2-fold. Targeting dCas9-P300 to the intronic regulatory elements increased DENND1A expression between 2.1-fold and 2.6-fold. The increase in DENND1A expression was statistically significant compared to the effect of the non-targeting gRNAs for the promoter and all four of the regulatory elements after Bonferroni correction for multiple hypothesis testing (Figure 5c, α < 0.05, t-test).

In H295R-dCas9-P300 cells, we observed a similar trend of increased DENND1A gene expression for on-target CRISPR perturbation compared to the effect of the non-targeting gRNAs. We observed a 2.2-fold increase in DENND1A expression when targeting dCas9-P300 to the DENND1A promoter. Targeting dCas9-P300 to the intronic regulatory elements increased DENND1A expression between 1.2-fold and 1.6-fold. The increase in DENND1A expression was statistically significant compared to the effect of the non-targeting gRNAs for the promoter and “element 3” of the regulatory elements after Bonferroni correction (Figure 5d, α < 0.05, t-test).

To test for off-target effects for genes in the DENND1A locus, we also measured gene expression changes for DENND1A flanking genes LHX2, CRB2, and STRBP. In both cell lines, we found CRB2 was not expressed and that expression of LHX2 and STRBP was not affected (Figure S22 and S23). We, therefore, inferred that the effects we observed were specific to DENND1A, and that genetic variation in the region likely contributes to PCOS via effects on DENND1A expression.

### Activation of PCOS-associated regulatory elements increased testosterone production in steroidogenic adrenal cells

Changes in gene expression levels might alter physiologically relevant phenotypes that could contribute to disease pathogenesis^66^. To test if endogenous overexpression of DENND1A could alter testosterone production in H295R cells, we overexpressed DENND1A by targeting dCas9-P300 to the DENND1A promoter or distal regulatory elements. We then measured the concentration of testosterone in the cell culture media four days later. Increasing DENND1A expression via activating the promoter caused a 3.2-fold increase in testosterone concentration, while activating three of the four distal regulatory elements individually increased testosterone concentration by between 1.7-fold and 2.2-fold (Figure 5e). The increases in testosterone concentration were statistically significant (α < 0.05, t-test). As a complementary analysis, we measured the rate of increase in testosterone concentration over the four days post transduction (Figure S24). Overall, the rate of change of testosterone concentration mirrored the levels measured after four days. Specifically, cells with increased DENND1A expression had substantially increased testosterone production between 2- and 4-days post-transduction compared to control-treated samples (Figure 5f). These results indicated that altered expression of endogenous DENND1A was sufficient to increase androgen biosynthesis in steroidogenic cells.

## Discussion

One of the central challenges of complex trait genetics is identifying the causal variants within GWAS susceptibility loci and determining their functional consequences. Here, we have fine mapped PCOS genetic associations to specific gene regulatory elements using a combination of high-throughput reporter assays and genetic association analyses. Specifically, we have mapped candidate regulatory elements by testing for the regulatory activity of millions of DNA fragments across 14 PCOS GWAS loci comprising of about 3 Mb of the human genome. We further demonstrated a scalable approach to fine map genetic variants within candidate regulatory elements. We identified novel PCOS-associated genetic variants by performing genetic association tests across genomic regions that we identified as candidate regulatory elements. Together, we demonstrated a generalizable strategy for identifying genetic variants within experimentally identified functional regulatory elements to fine map genetic association loci for complex genetic traits. As proof-of-concept of the strengths of this approach, we focused on DENND1A, a PCOS GWAS candidate gene reported to regulate androgen biosynthesis^22^. We showed that manipulating the epigenome of DENND1A-proximal regulatory elements caused increased DENND1A expression and, subsequently, increased androgen in human adrenal cells. These results extend previous studies identifying a role for DENND1A in testosterone production in theca cells, while also demonstrating specific gene regulatory elements wherein genetic variation can alter DENND1A expression. Our results demonstrate the advantage of combining high-throughput reporter assays, fine mapped genetic analyses, and targeted epigenome editing to discover novel gene regulatory mechanisms contributing to common human diseases.

The experimental approaches we used have several advantages and limitations. Because the targeted STARR-seq approach assayed fewer fragments, it was more amenable to be used in cell models that cannot be grown at large scales. The targeted approach also allowed us to test for regulatory activity outside context of genetic linkage^33^. Furthermore, the ability to capture natural genetic variation present in a pool of genomes allowed us to test for allele-specific regulatory activity across one locus in depth. It is understood that weak effects of non-coding variants contribute to a phenotype through coordinated regulation across several regulatory elements^67^. Thus, this approach allowed us to identify regulatory elements that contribute to an organismal phenotype through gene expression patterns.

A limitation in interpreting STARR-seq results was that the DNA fragments were not assayed in their chromatin context. For example, while the two cell lines used in this study have different steroidogenic properties, we observed very similar regulatory activities between them. That similarity is likely because STARR-seq models the regulatory activity of the DNA without influences from chromatin. In addition, we cannot show an effect on the target genes using a reporter assay. In this study we overcame those limitations using complementary CRISPR-based approaches.

Identifying the underlying mechanisms by which GWAS loci contribute to disease pathogenesis will be essential for translating these findings to benefit human health. The effect of regulatory elements and non-coding variants has been elucidated for several disease phenotypes. For example, one study identified a SNP that regulates SORT1 in a liver-specific manner within a GWAS risk locus for low-density lipoprotein cholesterol and myocardial infarction (MI)^66^. Another study focused on maternal hyperglycemia identified variants spanning multiple enhancers that have a coordinated effect on HKDC1 expression^68^. Other studies focused on post-GWAS functional analyses have used different methods, including statistical^49,69–75^ and experimental^76–81^ approaches to fine map GWAS signals and identify functional variants. Nevertheless, detailed cellular or molecular studies are often needed to connect the identified gene regulatory effects to a disease relevant phenotype^66,79^.

A challenge is the molecular follow up on putative causal genes, which is dependent on cell type, function of the genes and assays to measure the function of the gene with respect to the disease phenotype. PCOS, however, is particularly amenable to experimental perturbation since hormone responses are easy to model in cell systems and offer a potential for testing one of the main clinical phenotypes of PCOS. Our results extend the knowledge of non-coding genetic mechanisms of PCOS pathogenesis. Previous experimental studies characterized a highly conserved enhancer regulating FSHB expression in mouse pituitary cells^82,83^; and non-coding variants intronic to AMHR2, a receptor for anti-Müllerian hormone^84^. Previous statistical approaches have also nominated common and rare genetic variants altering the expression of DENND1A^24^, FSHB, ZFP36L2, ERBB3, RPS26, RAD50^85^ as potentially contributing to PCOS. Here, we add both a specific gene regulatory mechanism controlling DENND1A expression to that body of knowledge, while also demonstrating a general strategy for identifying analogous mechanisms for other PCOS genes.

The candidate regulatory elements that we identified can serve as a framework to identify functional non-coding regions that might contribute to PCOS risk by harboring causal variants. Our findings add to growing empirical evidence of regulatory regions contributing to complex traits^81,86–88^. We expect that future evaluation of the regulatory elements from this study will provide new insights into the mechanisms leading to PCOS phenotypes. Broadly, our results demonstrate a scalable approach to study disease-associated regulatory regions implicated not only in PCOS, but also in the pathogenesis of common, complex disorders in general.

## Methods

## Data Availability

All open chromatin regions and STARR-seq results for the studied regions are available via the Gene Expression Omnibus and a UCSC Genome Browser track hub (https://genome.ucsc.edu/s/laavatar/2024%2DPCOS%2DSS%2Dpaper).

## Acknowledgements

The Genotype-Tissue Expression (GTEx) Project was supported by the Office of the Director of the National Institutes of Health, and by NCI, NHGRI, NHLBI, NIDA, NIMH, and NINDS. The data used for the analyses described in this manuscript was obtained from the GTEx Portal on 7 June 2020.

## STARR Seq Assay Library Construction

### Selection of GWAS regions for targeted STARR-seq assays

To select PCOS-associated genomic regions for STARR-seq assays, we identified all genetic variants in linkage disequilibrium (LD, r^2^ > 0.8) with the 16 genetic variants that were most strongly associated with PCOS or its clinical phenotypes^9,11,13,14^. We then selected bacterial artificial chromosomes (BACs) and fosmids that encompassed all the identified genetic variants. We obtained a total of 18 BACs and 2 fosmids. All BACs and fosmid clones were sourced from BACPAC Genomics, Inc and the source of these clones is Children’s Hospital & Research Center at Oakland (CHRCO). The list of BACs and fosmids is detailed in Supplementary Table S1.

All BACs and fosmids were obtained as clones in E. coli. We propagated each bacterial clone in selective conditions. We isolated the BAC DNA using NucleoBond Xtra BAC (Machery-Nagel); and we isolated fosmid DNA using FosmidMAX (Lucigen), following manufacturer’s protocols. To validate that the BACs and fosmids were intact and covered the target region, we created Illumina high-throughput sequencing libraries from the isolated DNA using NEBNext Ultra II FS DNA Library Prep. We barcoded the sequencing library for each BAC or fosmid independently, and pooled the resulting libraries for sequencing. We sequenced the pooled libraries on an Illumina MiSeq instrument, and aligned to the human genome. For two of the 16 target regions, the BACs either recombined or the sequencing reads from the BAC aligned to a different genomic region suggesting contamination with another BAC. We removed those two regions from subsequent analysis. The BACs and fosmids for the remaining 14 target regions span ∼3 Mb of the human genome (Supplementary Table).

### STARR-seq reporter plasmid construction

To create STARR-seq assay libraries from the BACs and fosmids, we cloned sheared DNA from each BAC into the STARR-seq plasmid. We sheared each BAC or fosmid to ∼400 bp DNA fragments using a Covaris S220 sonication instrument. We then ligated custom universal adapters to the resulting DNA fragments using the NEBNext DNA Library Prep protocol (#E6040L) (Supplementary Table, S11 - SS_Adaptor_1 & SS_Adaptor_2). We amplified the adapted DNA fragments and added sequences for Gibson assembly into the STARR-seq plasmid using PCR. For the PCR, we used KAPA HiFi HotStart kit (Roche) and the primers TS2SS-F and TS2SS-R (Supplementary Table, S11). The PCR cycling conditions were: 98 °C for 30 s, followed by 10 cycles of 98 °C for 15 s, 64 °C for 30 s, 72 °C for 30 s, with a final extension at 72 °C for 5 min.

We cloned the fragment libraries into the STARR-seq ORI vector (Addgene#99296). To do so, we first linearized the plasmid using AgeI and SalI (NEB R3552L and NEB R3138L). We analyzed the digested plasmid on a 1% agarose gel, confirmed that the linear plasmid was the expected ∼3600 bp size, and isolated the linearized plasmid using either the QIAquick Gel Extraction Kit (#28704) or GeneJET Gel Extraction Kit (#K0691). We cloned the adapted and amplified DNA fragments from the BACs and fosmids into the linearised STARR-seq ORI vector using the NEBuilder HiFi DNA Assembly (#E2621) kit. We ethanol precipitated the products. To do so, we added 0.1X volume 3 M NaOAc and 2.5X volume cold 100% ethanol and stored the mixture at –20 °C overnight. We then pelleted the DNA via centrifugation at 16,000 RCF for 30 min at 4 °C. We washed the pellets with 5 ml cold 70% ethanol, and resuspended them in water. To amplify the resulting plasmid libraries, we electroporated into E. cloni 10G SUPREME Electrocompetent Cells following manufacturer protocol for optimal settings in 1.0 mm cuvette (10 μF, 600 Ohms, 1800 Volts). We grew the plasmids in individual 1 L volumes of LB with carbenicillin for antibiotic selection at 37 °C overnight. We isolated the resulting PCOS GWAS STARR-seq assay plasmids using NucleoBond PC 10000 EF (Machery-Nagel).

To make the final PCOS GWAS STARR-seq assay library, we pooled the individual BAC and fosmid STARR-seq plasmids in equimolar concentration. We validated the size of the plasmid library using the Agilent TapeStation, and quantified the resulting pool using Qubit (Invitrogen).

### PCOS GWAS STARR-seq assay library sequencing

To estimate the abundance of reads mapping to the regions selected in the assay library, we used Illumina high-throughput sequencing NextSeq 2000 with 50bp paired end sequencing protocol. To prepare the sequencing libraries, we first amplified the STARR-seq regions from a 20 ng pooled plasmid library using KAPA HiFi HotStart kit (Roche). The PCR cycling conditions were: 98 °C for 30 s, followed by 15 cycles of 98 °C for 15 s, 64 °C for 20 s, 72 °C for 30 s, with a final extension at 72 °C for 5 min using 208-F Index7 primers (Supplementary Table, S11). To isolate the final library, we used Axygen Spri Beads (AxyPrep™ Mag PCR Clean-Up Kit) beads at appropriate concentrations based on the manufacturer’s manual for an insert size of 400 bp.

#### Cell Culture protocol

We obtained NCI-H295R cells from ATCC. The cells were cultured in DMEM/F-12 medium (Gibco #21041025) supplemented with 2.5% Nu-Serum (Corning #355100) and 1% ITS+Premix (Corning #354352) and grown as a monolayer at 37 °C, 5%CO2. We validated testosterone produced by the cells stimulated with 10 µM forskolin using ELISA following manufacturer’s protocol (Cayman Chemicals #582701).

We obtained COV434 cells from ECACC (Sigma–Aldrich #07071909). The cells were cultured in DMEM (Gibco #11965092) supplemented with 2 mM Glutamine and 10% Foetal Bovine Serum (FBS) and grown as a monolayer at 37 °C, 5%CO2. We validated estradiol produced by these cells treated with 100 ng/mL follicle stimulating hormone (FSH) and 2.9 μg/mL androstenedione (A4) using ELISA (Cayman Chemicals #501890). All experiments were performed between passages 5 and 12.

#### Nucleofection Optimization

To transiently introduce the PCOS GWAS STARR-seq assay library into the cell lines, we used electroporation via the Lonza 4D-Nucleofector System. To optimize the electroporation settings for H295R and COV434, we used the Cell Line Optimization 4D-Nucleofector™ X Kit (Lonza #V4XC-9064) following manufacturer’s protocol. Based on this optimization, we chose SF-CM-138 for COV434 cells and SF-DN-100 for the H295R cells, with 2 μg of plasmid to every 1 million cells transfected.

#### Transfection of cells

To test the regulatory potential of PCOS GWAS targeted regions, we first transfected the PCOS GWAS STARR-seq library into both H295R and COV434 cells, and isolated and sequenced the resulting RNA. We isolated the RNA from the cells 6 hours post transfection. We transfected the PCOS GWAS STARR-seq plasmid library into H295R and COV434 based on the nucleofection optimization settings we described using SF Cell Line 4D-Nucleofector® LV Kit L (Lonza #V4LC-2002) following manufacturer’s protocol. All the experiments were performed in triplicate for each cell line. For each replicate for each of the cell lines, we used 50 million cells transfected with 100 μg of the PCOS GWAS STARR-seq plasmid library.

#### PCOS GWAS STARR-seq reporter library construction

To isolate the PCOS GWAS reporter RNA, we first isolated total RNA followed by enriching for cDNA produced from the PCOS GWAS STARR-seq library plasmid pool.

Six hours post transfection, we rinsed the cells with PBS and dissociated the cells using Trypsin-EDTA 0.25% (Life Technologies). We lysed the cell pellets using RLT buffer (Qiagen) with 2-mercaptoethanol (Sigma). We passed the lysates through a 18-gauge needle ten times and stored at −80 °C before RNA extraction.

### RNA extraction

We isolated total RNA using the Qiagen RNeasy Midi kit including the on-column DNaseI digestion step. We treated the isolated total RNA with 1 μL RNase Block (Agilent). We then isolated poly-A RNA using Dynabead Oligo-dT25 beads (Life Technologies) according to the manufacturer’s recommended protocol. We treated the poly-A RNA with DNase (TURBO DNase, Invitrogen) and 1 μL RNase Block at 37 °C for 30 min before halting the reaction with the DNase inactivation reagent. We then synthesized PCOS GWAS reporter cDNA by reverse transcription using Superscript III (800 U, Life Technologies) following manufacturer’s protocol and a STARR-seq specific primer (SSRT-UMI, Supplementary Table, S11.)

### PCOS GWAS STARR-seq reporter construction

Following synthesis, we treated the cDNA with RNaseA (Sigma) at 37 °C for 1 hour. We purified the PCOS GWAS reporter cDNA with SPRI beads (1.5X) and amplified using index-PCR primer and indexed PostSS-Index-5 primers (Supplementary Table, S11) to allow barcoding for sample multiplexing under the following conditions: 98 °C for 30 s, followed by 10-12 cycles of 98 °C for 10 s, 64 °C for 30 s, 72 °C for 30 s, with a final extension at 72 °C for 5 min. We split each sample into 7 individual PCR amplification reactions in this step. We determined the total number of cycles for amplification using a small portion of that sample in a qPCR protocol and estimating cycle number using 1/4^th^ the maximum plateau observed in the qPCR. We cleaned the amplified PCR products using SPRI beads (1.0X) and then validated the length distribution of the PCOS GWAS reporter library on Agilent tape station.

### PCOS GWAS STARR-seq reporter library sequencing

Final PCOS GWAS reporter libraries from each replicate experiment were pooled at equimolar 2nM concentrations. We sequenced the PCOS GWAS reporter libraries on Illumina NextSeq 2000 using 50bp PE sequencing.

#### Alignments and STARR-seq analysis

To estimate regulatory activity in the targeted PCOS GWAS regions, we used the abundance of the fragments expressed as RNA in the reporter library relative to their abundance in the assay library. We first aligned the PCOS GWAS assay library and the PCOS GWAS reporter library individually to the human genome (hg38) using bowtie2. We filtered reads with a quality score of Q>=30, and outside the centromeres and blacklisted regions. These reads were used for the downstream analysis. We used picardtools^89^ to mark and call duplicates. RPKM normalized STARR-seq read density was computed at single base pair resolution using deepTools^90^ utility bamCoverage. We used CRADLE^42^ package to correct biases and call peaks with the following options. We then estimated differential STARR-seq activity across the regions as fold change using DESeq^43^. We compared PCOS STARR-seq results from both COV434 and H295R cell lines to ATAC-Seq datasets generated for these cell lines. We also compared the peaks to the regulatory regions across ENCODE (V4) for both cell lines and primary tissues.

#### PCOS case-control variant association testing within candidate regulatory regions

To identify any association between genetic variants within functional STARR-seq regulatory elements and PCOS, we performed genetic association analyses. The selection criteria, clinical features and genotyping in 983 PCOS cases and 2951 controls from our previous stage 1 GWAS discovery cohort has been reported (Hayes, 2015). In brief, genotyping was performed using the Illumina OmniExpress (HumanOmniExpress-12v1_C) array^9^. Genotype imputation was performed using minimac4^91^ on the Michigan Imputation Server^92^ for phasing via Eagle^93^ using the TOPMED freeze 8 reference panel^94,95^. Variants were filtered to remove any SNPs with imputation quality (R^2^) less than 0.8 and restricted to STARR-seq regions of regulatory activity. PCOS association was tested within candidate regulatory regions. Single variant associations were carried out using PLINK^96^ on common variants (minor allele frequency [MAF]>1%) using logistic regression with PCOS as the outcome variable and age, BMI, and five principal components (PCs)^97^ as covariates. To control for false-positive discoveries, results were adjusted for Bonferroni correction thresholds. The number of PCOS cases was 983 rather than 984 reported in Hayes et al, 2015, because we updated the IBD exclusion criterion from 3rd degree relatives to 4th degree relatives resulting in the exclusion of 1 PCOS case.

#### Colocalization testing

To test for association between two datasets to identify likely causal SNP between two traits, we used a bayesian colocalization method^49^. For the PCOS-associated variants, we used the list of variants and its associated statistics from the above result for all variants with P < 0.3 (Table 3). We used the standard options for the colocalization testing. For the eQTL dataset, we used publicly available expression quantitative association data from the GTEx consortium GTEx Analysis V8 (dbGaP Accession phs000424.v8.p2, accessed on June 7, 2020). The GTEx dataset contains cis-eQTL data from ∼900 American donors of mostly European Ancestry (∼85%) across 49 tissues and of varying ages. We applied coloc, a Bayesian test for colocalization to identify the probability of a shared causal signal between the PCOS-regulatory element-associated variant and eQTL variants. We used the coloc.abf() function in the coloc R package with the default assignment of prior probabilities for a SNP being associated with each trait from the Coloc package. All analyses with a colocalization posterior probability (PP.4) > 0.3 using eQTL data from all tissues, adrenal tissue and ovarian tissue were reported in Table 4.

#### DENND1A-Enriched STARR Seq Assay Library Construction

To test for the allele specific regulatory activity of common variants, we modified the targeted STARR-seq assay to use a pool of 5 different human genomes instead of BACs and fosmids as described above. We selected genomes from individuals identified as female, and healthy from the 1000 Genomes project^61^. We used three individuals of European ancestry and two individuals of Han Chinese ancestry to be pooled into the targeted STARR-seq experiment to identify allele specific regulatory effects. The list of genomes used is listed in Supplementary Table.

### Enrichment for the targeted DENND1A region

We focused on variants present in the DENND1A locus, a region that spans the entire DENND1A gene and 100 kb upstream and downstream of the gene. For target enrichment of the DENND1A locus, we used targeting oligonucleotide probes. We first sheared each genome separately to ∼200 bp using Covaris (S220). We then used Agilent SureSelect Custom DNA Target Enrichment Probes to enrich the region around DENND1A (hg 38: chr9:123279654-124030107). We followed the Agilent SelectXT2 custom (Cat# 5190-4846) to enrich the target regions in each genome, however we modified the protocol at the adaptor ligation steps. We used a custom adaptor (SS_Adaptor) and amplified the resulting oligo fragments using TS2SS-F and TS2SS-R primers (Supplementary Table, S11).

### DENND1A locus STARR-seq reporter plasmid construction

To create the DENND1A STARR-seq assay libraries from the five genomes, we cloned the sheared and DENND1A-locus enriched DNA fragments. We cloned the amplified and enriched fragments into the linearised STARR-seq vector using NEBuilder HiFi DNA Assembly (#E2621). We ethanol precipitated the products. To do so, we added 0.1X volume 3 M NaOAc and 2.5X volume cold 100% ethanol, and stored the mixture at –20 °C overnight. We then pelleted the DNA via centrifugation at 16,000 RCF for 30 min at 4 °C. We washed the pellets with 5 ml cold 70% ethanol, and resuspended in water.

We then pooled the plasmids from each genome in equimolar concentrations. We amplified the pooled plasmids by transfecting the plasmids into E. cloni 10G Electrocompetent Cells following manufacturer protocol for optimal settings (1.0 mm cuvette, 10 μF, 600 Ohms, 1800 Volts). We subsequently isolated the plasmids using Qiagen Plasmid Kit, GigaPrep (Qiagen #12191), and quantified it using Qubit and validated the length of the pooled library on a 1% agarose gel. This purified, pooled plasmid is our DENND1A-locus STARR-seq library that was used for DENND1A-locus STARR-seq experiments.

### DENND1A locus STARR-seq assay library sequencing

To estimate the abundance of reads mapping to the variant loci selected in each assay library, we used Illumina high-throughput sequencing NextSeq 2000 with 50 bp paired end sequencing protocol. We sequenced 3 replicates of the DENND1A-locus STARR-seq assay library using the amplified the STARR-seq assay fragments from the pooled library using 208-F Index7 primers (Supplementary Table, S11).

### DENND1A locus enriched STARR-seq assay

To test for effects of variants in the targeted DENND1A locus, we transfected the DENND1A locus STARR-seq assay library into H295R cells, and isolated and sequenced the resulting RNA similar to the methods described previously. All experiments were performed in triplicate for each cell line. For each replicate for each of the cell lines, we used 70 million H295R cells transfected with 140 μg DENND1A locus STARR-seq assay library using the Lonza Nucleofector (setting SF-DN-100). We isolated the RNA from the cells 6 hours post transfection.

### DENND1A locus STARR-seq reporter library construction

To isolate the DENND1A locus reporter RNA, we first isolated total RNA followed by enriching for cDNA produced from the DENND1A-locus STARR-seq plasmid library. We used the same protocol as described for the PCOS GWAS reporter library construction. We pooled the DENND1A locus reporter libraries from each replicate at equimolar 2nM concentrations. We then sequenced the DENND1A locus reporter libraries on Illumina NextSeq 2000 using 75bp PE sequencing.

### Candidate regulatory variant identification

To identify variants that have allele specific regulatory activity, we compared the ratio of reads mapping to the alternate allele versus reference allele in each assay library and reporter library. If the ratio of reads mapping to alternate allele versus reference allele was higher in the reporter library compared to the assay library, that variant was called as having increased regulatory activity of the alternate allele.

To do so, we first obtained a list of the variants present in the DENND1A locus in the pool. We obtained the VCF for these samples from the 1000 Genomes Project^61^. We filtered the variants in the targeted DENND1A locus, to only include those SNPs present as heterogeneous within the pool of five genomes we used. The final list of ∼600 variants was then used for the regulatory variant analysis.

To compare reads mapping to each allele in both the reporter and assay libraries, we first aligned DENND1A locus enriched STARR-seq libraries (assay library and reporter library) individually aligned to the human genome (hg38) using WASP^98^ and bowtie2^99^. Reads with a quality score of Q>=30, and outside the centromeres and blacklisted regions were used for downstream analysis. We used picardtools to mark and call duplicates^89^. RPKM normalized STARR-seq read density was computed at single base pair resolution using deepTools utility bamCoverage^90^. We then assigned reads mapping to each variant for each sequenced sample using samtools mpileup^100^.

To estimate the regulatory effect of variants, we used BIRD^60^. BIRD is a bayesian statistical framework for analysis of regulatory variants and uses bayesian priors to identify allele-specific regulatory effects, and identifies variants that have a high probability of being a regulatory variant with an effect size, theta. We used the standard options for BIRD and set the regulatory effect threshold as 1.2.

#### ATAC-Seq

To identify the accessible chromatin within the H295R and COV434 cells, we performed ATAC-Seq^101^ in duplicate as described below.

We harvested 50,000 viable cells for each replicate. COV434 cells were additionally incubated in TURBO DNase (Invitrogen, #AM2238) for 1 hour at 37 °C. We then incubated the cells with 50 μL cold ATAC-RSB with 0.1% NP40, 0.1% Tween20, 0.01% Digitonin and incubated on ice for 3 minutes. We washed the cells with 1ml cold ATAC RSB with 0.1% Tween 20 and pelleted. We resuspended the cell pellets in the transposition mixture comprising of 25 μL TD buffer, 2.5 μL transposase, 16.5 μL PCS, 0.5 μL 1% digitonin, 0.5 μL 10% Tween 20 and 5 μL H2O and incubated in a thermomixer at 37 °C for 30 minutes. We cleaned up the DNA using MinElute Reaction Cleanup Kit (Qiagen, #28204). We amplified the resulting DNA using an ATAC-Universal primer and an ATAC-barcode primer (Supplementary Table, S11) and cleaned it using SPRI beads. We sequenced the ATAC seq libraries on Illumina NextSeq 550, 50 bp PE sequencing at the Duke Genomics core.

##### ATAC-seq preprocessing and alignment

ATAC-seq libraries for H295R and COV434 cell lines were individually aligned to the human genome (hg38). Each cell line had 2 biological replicates, and >40 million reads were generated per sample. Sequencing data quality was assessed with FastQC, and adapters were trimmed with Trimmomatic. Trimmed reads were aligned to the GRCh38 genome using Bowtie^99^ reporting only alignments having no more than two mismatches, discarding multi-mapping reads(-v 2 --best --strata -m 1). Reads mapping to the ENCODE hg38 blacklisted regions (https://www.encodeproject.org/files/ENCFF356LFX; manually curated regions with anomalous signal across multiple genomic assays and cell types) were removed using bedtools2 intersect^102^ (v2.25.0). Properly paired reads were then filtered to exclude presumed PCR duplicates using Picard MarkDuplicates (v1.130; http://broadinstitute.github.io/picard/). Reads were then used to generate reads per million (RPM) counts of bigWig files for visualization using deeptools bamCoverage^103^ (v3.0.1). Peaks were called using MACS2 with an FDR cutoff 0.1. We used the ENCODE ATAC-Seq standards for analysing the dataset we generated. We generated Transcription Start Site enrichment values using GRCh38 Refseq TSS annotation and used the cutoff of >7 for high quality data (Figure S8).

#### Generating cell lines for CRISPRa perturbation studies

##### GuideRNA (gRNA) design and gRNA plasmid synthesis

Four candidate regulatory elements were identified from the targeted STARR-seq results with coordinates listed in Table 9. The regions were selected based on STARR-seq effect, chromatin accessibility and ability to design guides considering genomic sequence and PAM restrictions.

To design the guide oligos, we used Guidescan2^104^, with “specificity” filter > 0.2. We had a total of 21 gRNAs, across four candidate regulatory elements and DENND1A promoter region (Supplementary Table), with each regulatory element comprising of 5-7 guides targeting that element. For the negative control, we designed a set of five guides that did not have any targets in the human genome. Each gRNA oligo was synthesized as individual oligos that were then processed as described below to make pooled gRNA plasmids.

To make the gRNA plasmids, we followed the outline of the CROP-Seq protocol^105^. First, we prepared the gRNA plasmid backbone by digesting CROPseq-Guide-Puro plasmid from Addgene (#86708) using BsmBI. We ran the digested product on 1% agarose gel, and we purified the 8.3 kb fragment using GeneJET Gel Extraction Kit (#K0691). To prepare the gRNA oligos for insertion into the plasmid, for each gRNA oligo synthesized, we first converted it to a double stranded oligo using Primers ssds-F and ssds-R

(Supplementary Table, S11). We then cloned each double-stranded gRNA oligo into the digested CROPseq-Guide-Puro vector using NEBuilder HiFi DNA Assembly (#E2621) kit. The plasmid products were purified with QIAquick PCR Purification Kit (Qiagen #28104). To make the pooled plasmids, we pooled (equimolar) each plasmid product for each regulatory element, or promoter region, or negative control. To amplify the plasmid pools, we electroporated each pool into Lucigen Endura Cells (Lucigen #60242-2) following manufacturer protocol for optimal settings in 1.0 mm cuvette (25 μF, 200 Ω, 1.5 kV). We grew the plasmids in individual 25 mL volumes of LB with carbenicillin for antibiotic selection at 37 °C overnight and isolated the gRNA plasmid pools using Qiagen Midi Prep (Qiagen #12143) following manufacturer’s protocols. Each purified plasmid pool was then used to prepare lentiviral particles.

### Lentivirus production

To test the target gene of the identified STARR-seq regulatory elements, we used CRISPRa to perturb the selected candidate regulatory elements. First, we designed a stable cell line expressing a Cas protein. To do so, we used a catalytically inactive Cas9 (dCas9) fused with the P300 domain of histone acetyltransferase (dCas9-P300). This dCas9-p300 can act as a transcriptional activator when combined with targeting guide RNA^37^.

To make stable dCas9-P300 cell lines, we generated lentivirus expressing dCas9-p300. Briefly, we combined the following plasmids: dCas9-p300 (Addgene #83889), psMD2.G (Addgene #12259) and psPAX2 (Addgene #12260) with Lipofectamine 3000 (Invitrogen #L3000001) and lipofected into HEK293T cells (ATCC #CRL-3216™) according to the manufacturer’s protocol. After 14 to 20 hours, transfection media was exchanged with fresh media. We then harvested viral supernatant at 24 and 48 hours post lipofection. We concentrated the viral supernatant at 1/100x using LentiX Concentrator (Clontech #631232) following the manufacturer’s protocols.

To produce lentivirus for individual gRNAs, we transfected HEK293T cells with an equimolar pool of gRNA plasmids for each regulatory element, psPAX2, and pMD2.G using Lipofectamine 3000 following the manufacturer’s instructions. We harvested media containing the produced lentivirus at 24 and 48 hours later and concentrated the viral supernatant at 1/100x using LentiX Concentrator (Clontech #631232) following the manufacturer’s protocols.

HEK293T cell line with stable dCas9-P300 expression:

We received HEK293T-dCas9-P300 cell line^37^ from Dr. Charles Gersbach. We followed the published culture and growth conditions for 293T cells.

### Generating stable H295R-dCas9-P300 cell line

To make stable H295R cells expressing dCas9-P300, we transduced the concentrated lentiviral particles containing dCas9-p300 into H295R cells with a multiplicity of infection of 5.0 using 6 μg/ml of polybrene (EMD Millipore Corporation #TR-1003-G). Additionally, we selected for the transduced cells using 0.5 μg/mL of puromycin (Gibco #A1113803). We confirmed the expression of dCas9-p300 in H295R cells using qRT-PCR.

### Transduction of gRNA into dCas9-P300 expressing cell lines

To test the effect of P300 on the targeted regulatory elements, we transduced each lentiviral pool for the regulatory elements, DENND1A promoter region and negative control in two cell lines (HEK293T and H295R) with stable dCas9-P300 expression. We transduced the cells during seeding in a 12-well or 6-well plate supplemented with 6 μg/ml of polybrene for H295R cells and 4 μg/mL of polybrene for HEK293T cells across 6 replicates for each pool (EMD Millipore Corporation #TR-1003-G). We changed the media on the cells 24 hours after transduction.

#### RNA isolation and qRT-PCR to measure gene expression levels

To measure any changes in gene expression levels due to the CRISPRa perturbation, we used qRT-PCR. First, we harvested RNA from each replicate 4 days post transduction with the gRNA lentivirus pool using RNeasy Mini Kit (Qiagen #4004) following manufacturer’s protocol including the DNase treatment. We quantified the RNA using Qubit (Invitrogen) and used 500 ng of RNA for each sample for subsequent cDNA synthesis. For the cDNA synthesis, we used Superscript III (800 U, Life Technologies) with Oligo dT primers following manufacturer’s protocol (Thermo Fisher #18418012). Following cDNA synthesis, we performed qRT-PCR using that cDNA, TaqMan™ Fast Advanced Master Mix for qPCR (Thermo Fisher #4444556), and TaqMan™ Gene Expression Assays (for the genes DENND1A, CRB2, LHX2 and STRBP and GAPDH). The qPCR analysis was performed using the 2^-ΔΔCT^ method in R, using GAPDH as the internal control. All the fold change is reported as log(2^-ΔΔCT^) compared to the negative (non-targeting gRNA) control. Each sample was measured in triplicate for the qRT-PCR.

#### ELISA for measuring testosterone production

To measure changes in testosterone production, we collected the supernatant from the gRNA pool transduced H295R cells two- and four-days post transduction. First, we diluted the supernatant 10-fold. Then, we measured the amount of testosterone produced using ELISA (Cayman Chemicals #582701) according to the manufacturer’s protocols using the given standard. All samples were measured in triplicate. The absorbance of the compound was measured at 405-420 nm using the GloMax Discover System (Promega). Fold-change reported is based on the negative (non-targeting gRNA) control.

## Supplementary figures

S1: Number of chromatin accessible regions within the selected coordinates for targeted STARR-seq using BAC

S2: Fragment length distribution in PCOS-GWAS loci targeted STARR-seq: Length of the fragments (bp) present in the assay (light blue) and reporter (dark blue) libraries.

S3: PCOS GWAS STARR-seq library complexity: The number of expected unique fragments of the reporter libraries for H295R (purple) and COV434 cells (teal) estimated using preseq. At 3×10e^9^ total reads, expected unique reads would be 2.8×10e^8^ for H295R and 3.2×10e8 for COV434 respectively.

S4 Correlation across assay libraries for STARR-seq: Pearson correlation for the assay (input) libraries for targeted STARR-seq for H295R cells (top) and COV434 cells (bottom). Scatterplots comparing log_10_-transformed CRADLE corrected read counts across chromatin accessible regions across three replicates of STARR-seq assay libraries.

S5: Correlation across assay and reporter libraries for STARR-seq: Scatterplots comparing log_2_-transformed reporter libraries over assay libraries as log_2_-fold changes for H295R cells (top) and COV434 cells (bottom) after CRADLE correction. Assay library replicates assigned randomly to reporter library replicates 1, 2 and 3.

S6: PCA plots demonstrating successful clustering of each replicate of assay and reporter libraries for STARR-seq for both H295R (purple) and COV434 (teal) cells.

S7: Scatterplot of STARR-seq effect sizes of regulatory elements shared between COV434 and H295R cells based on Jaccard index > 0.5. The effect sizes are measured on CRADLE-corrected and CRADLE-called regulatory elements using DESeq2.

S8: PCA plots demonstrating successful clustering of each replicate of ATAC-seq for both H295R (purple) and COV434 (teal) cells.

S9: Venn diagram indicating the shared and unique accessible chromatin regions in H295R and COV434 based on ATAC-seq. ATAC-seq regions were called using MACS2 with FDR < 0.1.

S10: Transcription start site (TSS) enrichment profiles for ATAC-seq of H295R (top) and COV434 (bottom) cells. The enrichment scores for both cell lines were > 7.

S11: Scatter plot of effect sizes measured using DESeq for both COV434 and H295R cells. The regions were selected as chromatin accessible regions that also were called in CRADLE as a peak and had regulatory activity measured in at least one cell line.

S12: Boxplots of effect size of candidate regulatory elements corresponding to DHS overlap: Candidate regulatory elements in H295R cells (purple) and COV434 cells (teal) with increasing activity correspond to regions with increased evidence of functionality using ENCODE DHS data.

S13: (a). Aggregate profile plots of chromatin accessibility based on ENCODE DNaseI Hypersensitive sites (DHS) centred on the active candidate regulatory elements across 400 bp windows for both cell lines (H295R in purple, COV434 in teal). Control regions (grey) are randomly generated genomic regions that are chromosome-, length- and GC-matched to the STARR-seq elements. (b). Aggregate profile plots of conservation score (phastcons) across active STARR-seq regulatory elements in H295R (purple) and COV434 (teal). The control regions are in grey and are random GC- and length-matched to the STARR-seq regions.

S14: Aggregate profile plots of conservation score (phastcons) centered on the ATAC-seq peak. ATAC-seq regions shared between two cell lines (H295R and COV434) are shown in blue, cell-specific ATAC-seq regions are in yellow and random GC- and length-matched control regions in cyan.

S15: Genome browser overview of the regulatory elements identified in DENND1A locus. Tracks shown : H295R STARR-seq (purple), COV434 STARR-seq (teal), H295R and COV434 ATAC-seq (grey), ENCODE cCRE : yellow and orange : distal and prximal Enhancer like elements, ref : promoer like elements. SNPs identified within the candidate regulatory elements are shown in blue vertical bars. The linkage disequilibrium heatmap was generated using EUR data from the 1000 Genomes Project

S16: DENND1A enriched STARR-seq library complexity: The number of expected unique fragments of the reporter libraries in H295R cells estimated using preseq. At 1×10e^8^ total reads, expected unique reads would be ∼0.7e^6^ unique reads averaging at 140x coverage per base pair.

S17: Fragment length distribution in DENND1A-enriched STARR-seq: Length of the fragments (bp) present in the assay (light blue) and reporter (dark blue) libraries.

S18: Correlation across DENND1A-enriched STARR-seq libraries: Pearson correlation for the assay (input) libraries in H295R cells (top). Scatterplots comparing log_10-_transformed read counts across chromatin accessible regions across three replicates of assay libraries. Scatterplots comparing log_2_-transformed reporter libraries over assay libraries as fold changes in H295R cells (bottom).

S19: Distribution of log_2_-transformed effect sizes for variants in DENND1A locus estimated using BIRD model: The variants highlighted in blue have a posterior probability of that variant being a regulatory variant > 0.9.

S20: Regulatory variants estimated using BIRD overlap known region of functional evidence including ENCODE cCRE - distal enhancer-like elements (dELS), promoter-like elements (PLS), proximal enhancer-like elements (pELS), CTCF binding sites (CTCF) and DNaseI hypersensitive sites (DHS).

S21: DENND1A locus (representative genome browser screenshot, ∼550 kb total length of the gene) with candidate regulatory elements for CRISPR-dCas9-p300 perturbation (in red). Each STARR-seq track is reported as assay (input) subtracted reporter (output) libraries.

S22: Log-fold change of LHX expression (left) and STRBP expression (right) with GAPDH as control for HEK293T cells stably expressing dCas9-p300. A set of 3 non-targeting control guides were designed to not target any part of the human genome as control cell population.

S23: Log-fold change of LHX expression (left) and STRBP expression (right) with GAPDH as control for H295R cells stably expressing dCas9-p300. A set of 3 non-targeting control guides were designed to not target any part of the human genome as control cell population.

S24: Testosterone concentration (pg/ml) measured in the cell media 2-days post transduction in H295R-dCas9-p300 cells targeted with the specific guide RNA.

