## Supplemental Tables and FIgures for "Gene regulatory activity associated with PCOS revealed *DENND1A*-dependent testosterone production"

Genomic coordinates (hg38)

| | BAC/fosmid ID | PCOS GWAS locus | chr | Start | End | In final PCOS STARR-seq library? |
|---|---|---|---|---|---|---|
| 1 | RP11-352H17 | LHCGR | chr2 | 48567960 | 48773736 | Yes |
| 2 | RP11-825O6 | FSHR | chr2 | 48947410 | 49120420 | Yes |
| 3 | RP11-17L5 | THADA | chr2 | 43201351 | 43356486 | Yes |
| 4 | RP11-831F19 | THADA | chr2 | 43348866 | 43512262 | Yes |
| 5 | RP11-677O13 | RAD50/IRF1 | chr5 | 132440666 | 132624332 | Yes |
| 6 | RP11-118D21 | GATA4/NEIL2 | chr8 | 11702874 | 11884058 | Yes |
| 7 | RP11-959L16 | DENND1A | chr9 | 123670418 | 123860422 | Yes |
| 8 | RP11-885N4 | AOPEP | chr9 | 94874999 | 95023646 | Yes |
| 9 | Rp11 640G3 | YAP1 | chr11 | 102141887 | 102311386 | Yes |
| 10 | RP11-1023M3 | RAB5B/SUOX | chr12 | 55873705 | 56034531 | Yes |
| 11 | RP11-1030M2 | KRR1 | chr12 | 75372807 | 75564639 | Yes |
| 12 | RP11-462A13 | HMGA2 | chr12 | 65796499 | 65964907 | Yes |
| 13 | RP11-261N13 | TOX3 | chr16 | 52296365 | 52442860 | Yes |
| 14 | RP11-159F20 | SUMO1P1 | chr20 | 53800706 | 53974652 | Yes |
| 15 | CH17-438B10 | FSHB | chr11 | 30024172 | 30225507 | Yes |
| 16 | CH17-267K24 | FSHB | chr11 | 30272821 | 30482841 | Yes |
| 17 | WI2-1214M9 | FSHB | chr11 | 30239629 | 30280497 | Yes |
| 18 | WI2-2586K13 | FSHB | chr11 | 30211547 | 30250398 | Yes |
| 19 | CTD-2564J11 | INSR | chr19 | 7101551 | 7273107 | No |
| 20 | RP11-367A1 | ERBB4 | chr2 | 212363496 | 212570765 | No |

Sum of megabases covered **2902898**

S1: Selected STARR-seq regions that have an accessible chromatin regions (ATAC-seq)

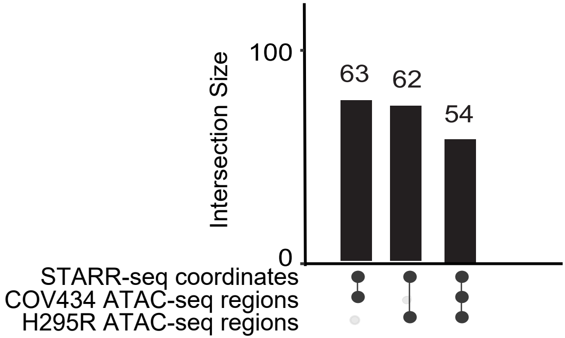

S2: Distribution of lengths of fragments in assay and reporter libraries in targeted STARR-seq experiments

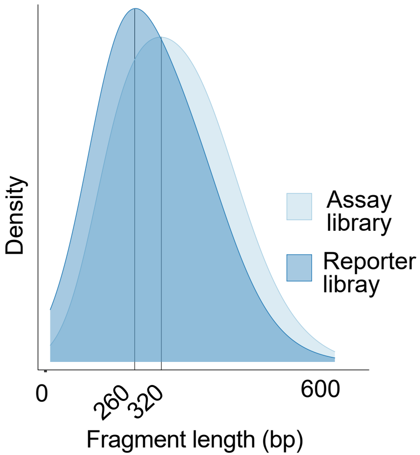

S3: Pre-seq complexity extrapolation plot for targeted STARR-seq libraries

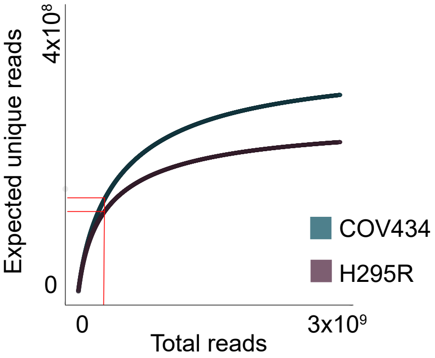

S4: Pearson correlation for STARR-seq assay libraries

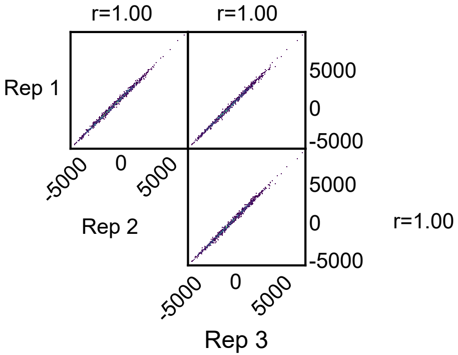


H295R assay library replicates

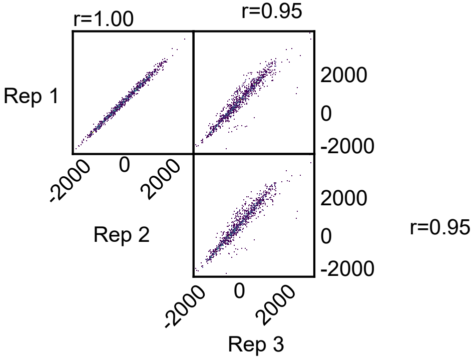


COV434 assay library replicates

S5: Pearson correlation of log2 (fold change) of reporter libraries to assay libraries

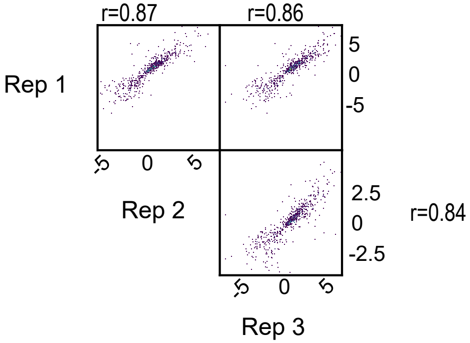


H295R

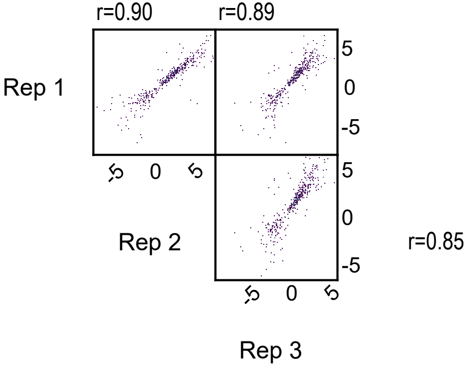


COV434

### S6: PCA for STARR-seq reporter and assay libraries

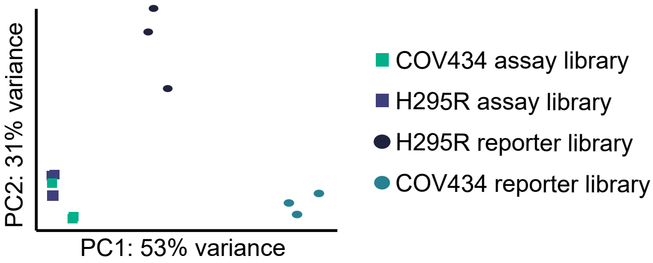

### S7: $Log_2$ (fold change) of regulatory elements with 50% or higher overlap

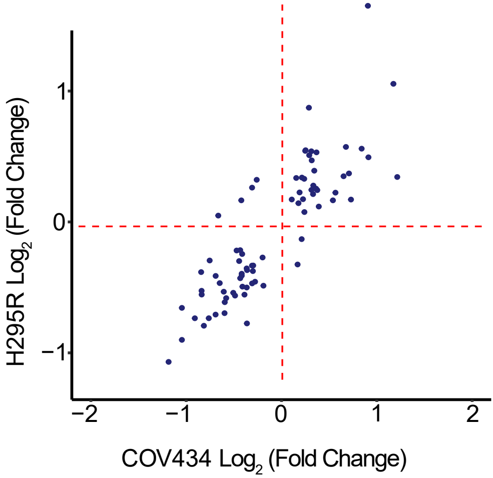

S8: PCA of ATAC-seq libraries for COV434 and H295R

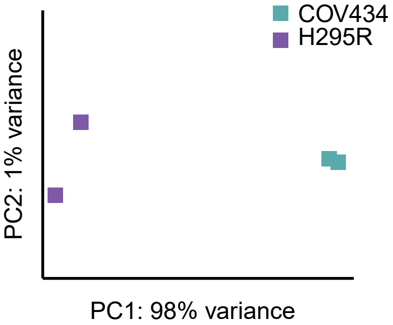

S9: Accessible chromatin regions in H295R and COV434 using ATAC-seq

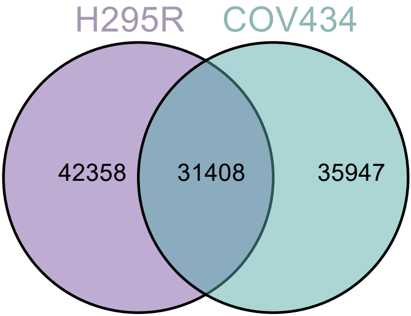

### S10: Transcription Start Site enrichment profiles for ATAC-seq of H295R cells (top) and COV434 cells (bottom)

H295R

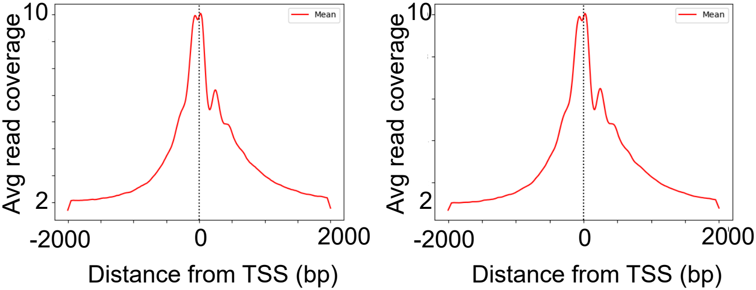


COV434

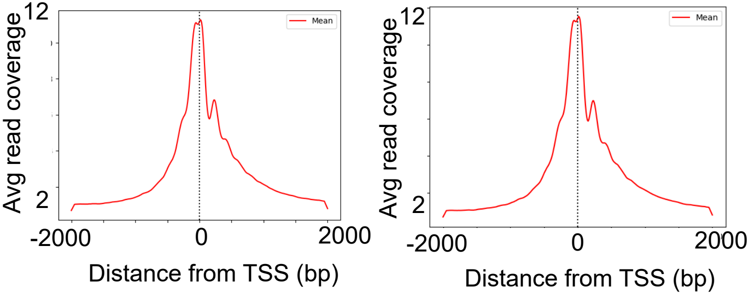

### S11: Effect sizes across open chromatin regions

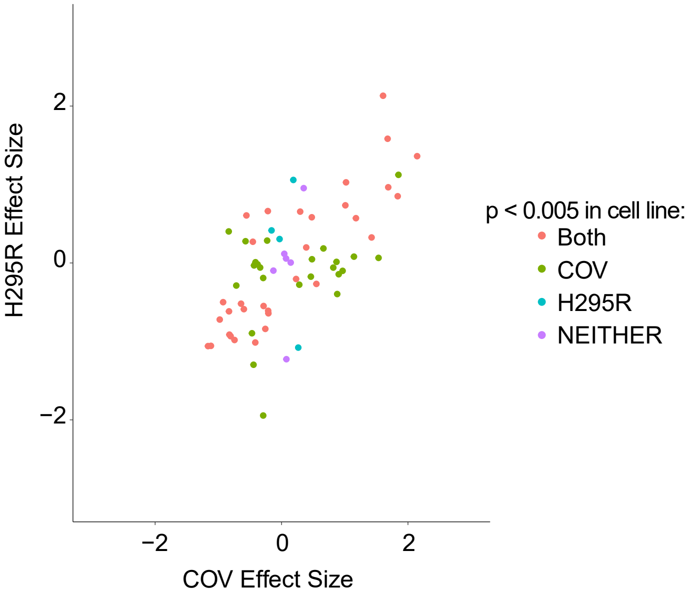

STARR-seq regulatory elements that overlap accesssible chromatin in H295R cells

| chr | start | end |
|---|---|---|
| chr11 | 30251372 | 30251872 |
| chr11 | 30322922 | 30323622 |
| chr11 | 102268237 | 102268887 |
| chr11 | 102287869 | 102288362 |
| chr12 | 55926915 | 55927405 |
| chr12 | 55959955 | 55960605 |
| chr12 | 55974205 | 55974405 |
| chr12 | 55997605 | 55997905 |
| chr12 | 56007605 | 56008405 |
| chr12 | 65826349 | 65826543 |
| chr12 | 65875749 | 65876049 |
| chr12 | 65891999 | 65892499 |
| chr12 | 65925793 | 65926477 |
| chr12 | 65948449 | 65949199 |
| chr12 | 75389507 | 75389857 |
| chr12 | 75390957 | 75391657 |
| chr12 | 75433095 | 75433295 |
| chr12 | 75480507 | 75480807 |
| chr12 | 75511407 | 75512042 |
| chr2 | 43332751 | 43333051 |
| chr2 | 43333951 | 43334401 |
| chr2 | 48601105 | 48601604 |
| chr5 | 132459216 | 132459916 |
| chr5 | 132491216 | 132491366 |
| chr5 | 132496356 | 132496456 |
| chr5 | 132506396 | 132506596 |
| chr5 | 132526216 | 132526916 |
| chr5 | 132556616 | 132557316 |
| chr8 | 11736452 | 11736863 |
| chr8 | 11769452 | 11769924 |
| chr8 | 11802474 | 11802774 |
| chr8 | 11853774 | 11854374 |
| chr8 | 11864313 | 11864463 |
| chr8 | 11867760 | 11868259 |
| chr9 | 94937135 | 94937879 |
| chr9 | 95003963 | 95004113 |
| chr9 | 95004513 | 95004889 |
| chr9 | 95004985 | 95005747 |
| chr9 | 123752924 | 123753173 |

STARR-seq regulatory elements that overlap accesssible chromatin in COV434 cells

| chr | start | end |
|---|---|---|
| chr11 | 102224081 | 102224231 |
| chr11 | 102268437 | 102268887 |
| chr11 | 102269237 | 102269337 |
| chr11 | 102294063 | 102294213 |
| chr12 | 55956145 | 55956445 |
| chr12 | 56007736 | 56007936 |
| chr12 | 56029670 | 56030011 |
| chr12 | 65858899 | 65859199 |
| chr12 | 65877119 | 65877241 |
| chr12 | 65896199 | 65896399 |
| chr12 | 65948599 | 65949299 |
| chr12 | 75390450 | 75390897 |
| chr2 | 43222151 | 43222251 |
| chr2 | 43226001 | 43226351 |
| chr2 | 43227101 | 43227451 |
| chr2 | 43240401 | 43240751 |
| chr2 | 48600960 | 48601410 |
| chr20 | 53805956 | 53806106 |
| chr20 | 53828006 | 53828606 |
| chr20 | 53829206 | 53829806 |
| chr20 | 53890756 | 53891006 |
| chr20 | 53900406 | 53900906 |
| chr20 | 53901156 | 53901356 |
| chr20 | 53915306 | 53915656 |
| chr20 | 53915806 | 53915906 |
| chr20 | 53942074 | 53942672 |
| chr5 | 132474574 | 132475782 |
| chr5 | 132477690 | 132477990 |
| chr5 | 132495899 | 132496299 |
| chr5 | 132496349 | 132496499 |
| chr5 | 132556745 | 132557245 |
| chr8 | 11765310 | 11765739 |
| chr8 | 11802224 | 11802410 |
| chr8 | 11845674 | 11846024 |
| chr8 | 11853724 | 11854524 |
| chr8 | 11874774 | 11875374 |
| chr9 | 95008207 | 95008503 |

S12: Effect sizes of candidate regulatory elements with ENCODE DHS overlap

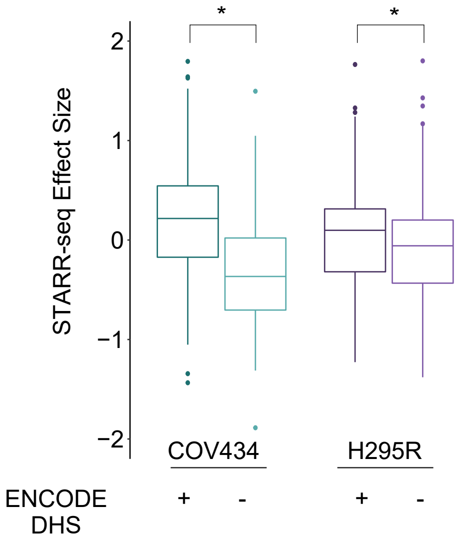

### S13: Active STARR-seq regulatory elements correspond to regulatory activity in multiple cell types and have increased conservation score

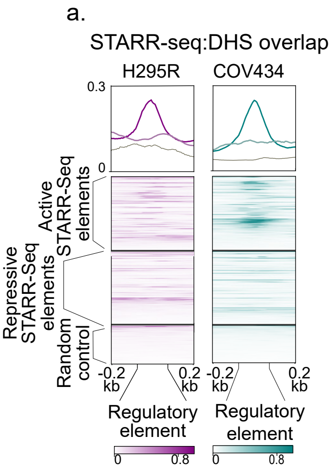


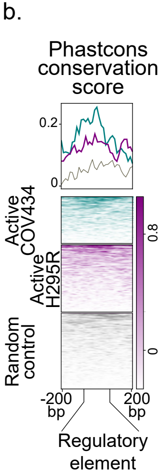

S14: Enrichment profile for ATAC-seq regions (in H295R and COV434) and phastcons conservation sore

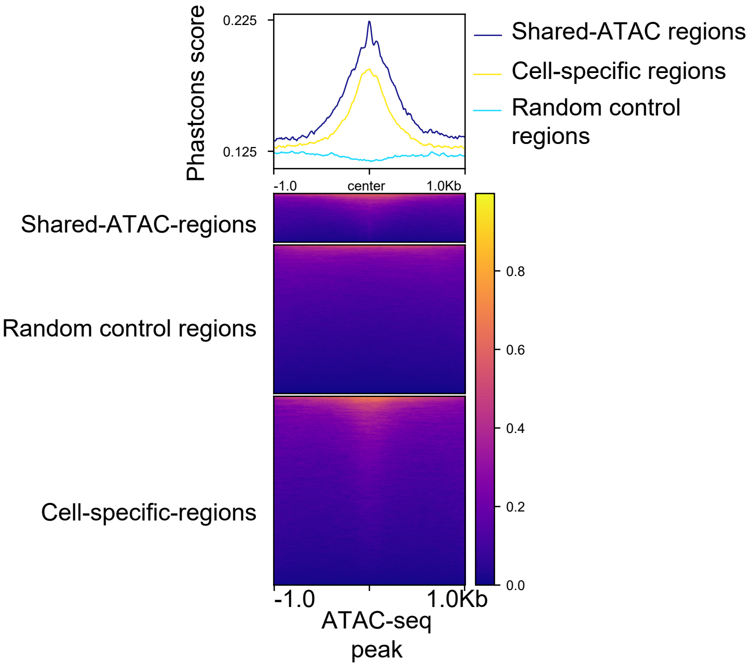

| SNP | SNP.AllTissue | Gene(s) | PP.in.OvaryTissue | PP.in.AdrenalTissue |
|---|---|---|---|---|
| **chr8:11769705** | **1** | **NEIL2** | 0.8792028 | |
| **chr12:55999553** | **1** | **RPS26/RAB5B/SUOX** | | 0.26656075 |
| **chr20:53949370** | **0.8613405** | **BCAS1** | | |
| **chr9:123707451** | **0.84471762** | **DENND1A** | | |
| **chr2:48730442** | **0.761973** | **LHCGR** | | |
| **chr11:30341402** | **0.7054277** | **ARL14EP/FSHB** | 0.16 | 0.34196682 |
| **chr9:94907391** | **0.6852266** | **C9orf3/AOPEP** | | |
| chr16:52298257 | 0.4462105 | | | |
| chr16:52305423 | 0.4455078 | | | |
| chr9:94909887 | 0.3143894 | | | |
| chr2:48740334 | 0.2375562 | | | |
| chr11:30323214 | 0.145648 | | 0.14 | 0.24537309 |
| chr11:30323252 | 0.145648 | | 0.14 | 0.24537309 |
| chr20:53802722 | 0.1386595 | | | |
| chr5:132492133 | 0.13864575 | | 0.07 | |
| chr5:132486441 | 0.12182875 | | 0.07 | |
| chr5:132487495 | 0.12182875 | | 0.07 | |
| chr5:132486380 | 0.11976187 | | 0.06 | |
| chr5:132486532 | 0.11976187 | | 0.06 | |
| chr5:132485496 | 0.11405538 | | 0.06 | |
| chr5:132485554 | 0.11405538 | | 0.06 | |
| chr9:123858278 | 0.0995106 | | | |
| chr16:52419655 | 0.0926816 | **TOX3** | 1 | 1 |
| chr5:132479665 | 0.07690871 | | 0.06 | |
| chr5:132496074 | 0.06264267 | | 0.09 | |
| chr9:123810883 | 0.03777879 | | | |

### S15: Genome browser map of regulatory elements identified in DENND1A locus including LD map

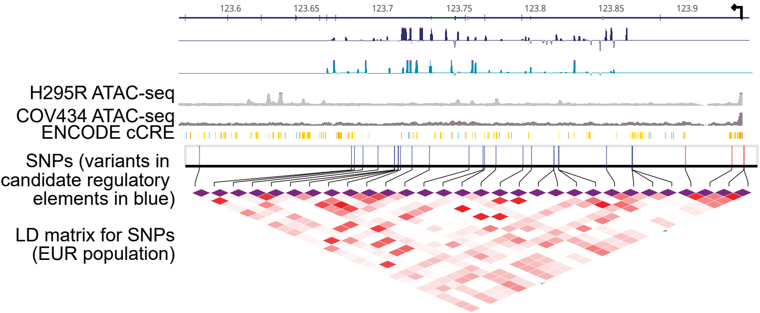

| 1KGP names | Ancestry |
|---|---|
| NA12616 | CEPH |
| NA06989 | CEPH |
| NA12832 | CEPH |
| HG00410 | CHS |
| HG00452 | CHS |

S16: Pre-seq complexity extrapolation plot for DENND1A-enriched STARR-seq

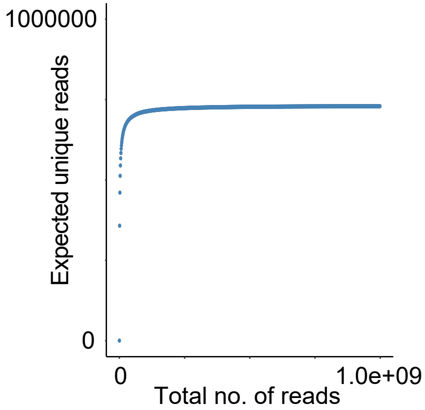

S17: Distribution of lengths of fragments in assay and reporter libraries in DENND1A-enriched STARR-seq experiments

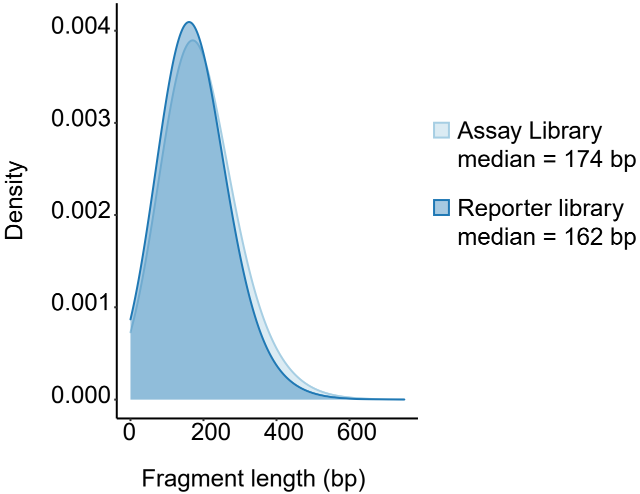

S18: Assay and log(fold change) of reporter library correlation plot for DENND1A-enriched STARR-seq

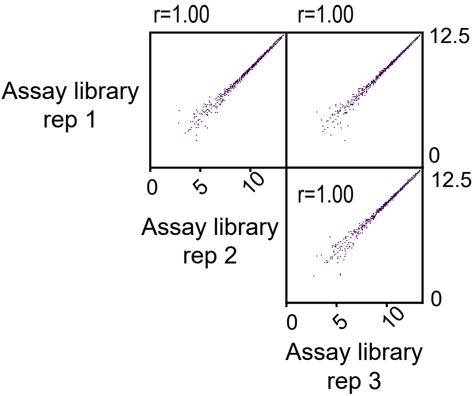


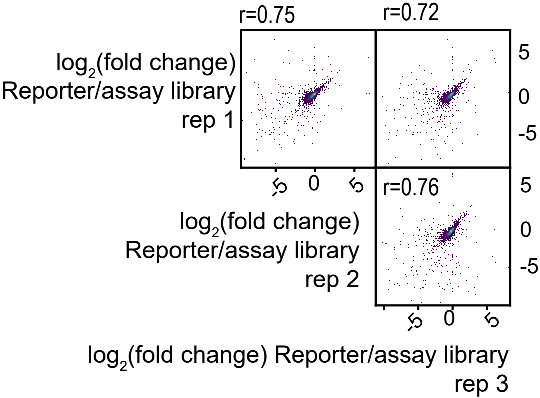

S19: Distribution of effect sizes caluclated for variants in DENND1A locus

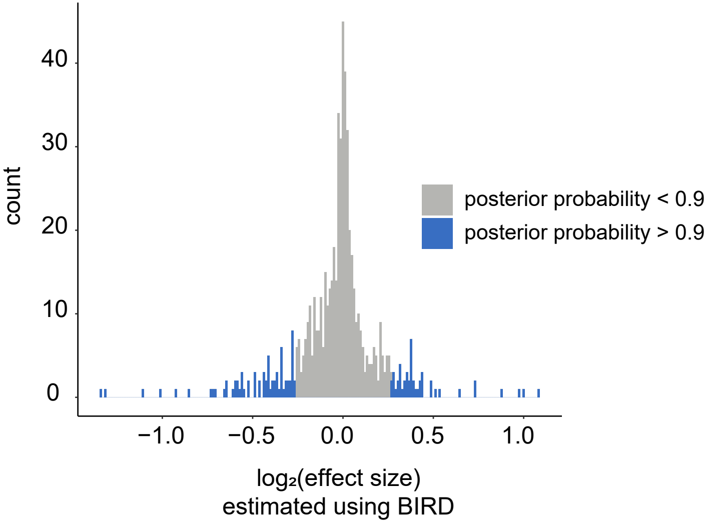

### S20: Overlap of top candidate regulatory variants with ENCODE genomic datasets

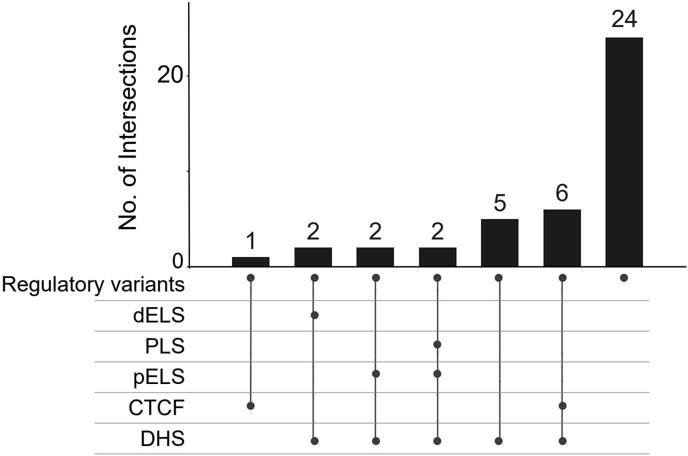

S21: Genome browser representation of DENND1A locus with regulatory elements targeted for perturbation

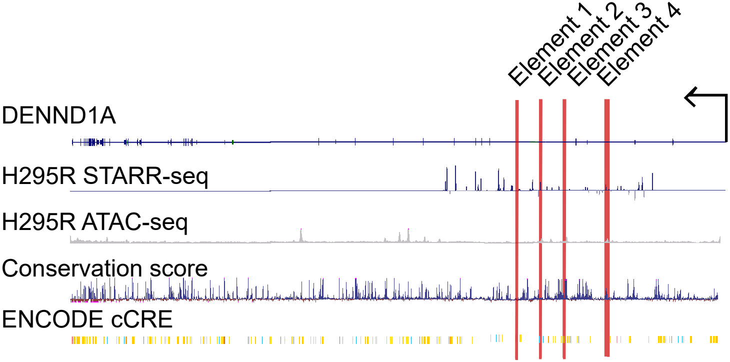

| | Genomic coordinates (hg38) | | |
|---|---|---|---|
| **gRNA pool name** | **chromosome** | **start** | **end** |
| Element 1 | chr9 | 123704717 | 123705190 |
| Element 2 | chr9 | 123724455 | 123724767 |
| Element 3 | chr9 | 123752897 | 123753207 |
| Element 4 | chr9 | 123770099 | 123770523 |
| Promoter | chr9 | 123929973 | 123930568 |

S22: Gene expression for STRBP and LHX (GAPDH control) via RT-qPCR in HEK293T-dCas9-p300

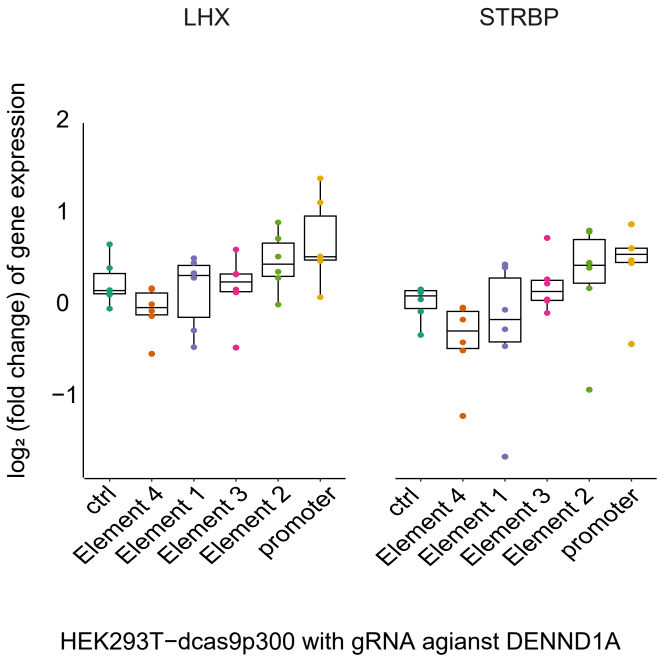

S23: Gene expression for STRBP and LHX (GAPDH control) via RT-qPCR in H295R-dCas9-p300

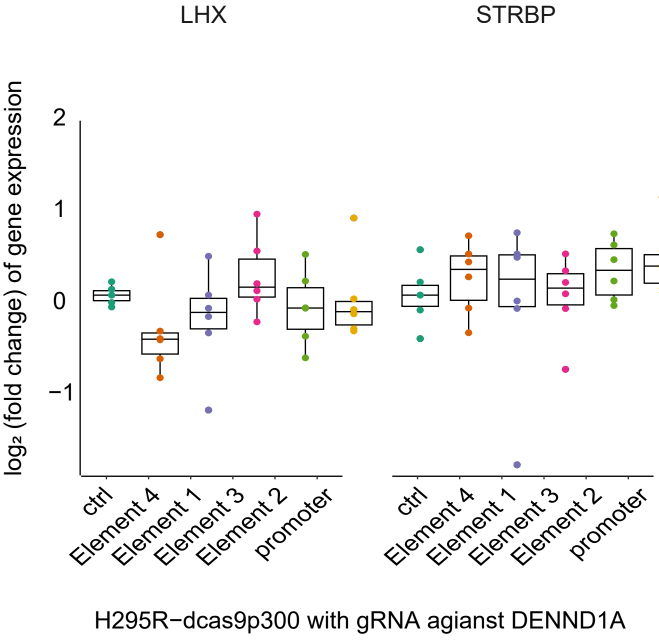

S24: Testosterone concentartion via ELISA in H295R-dCas9-p300, 2 days-post-transduction

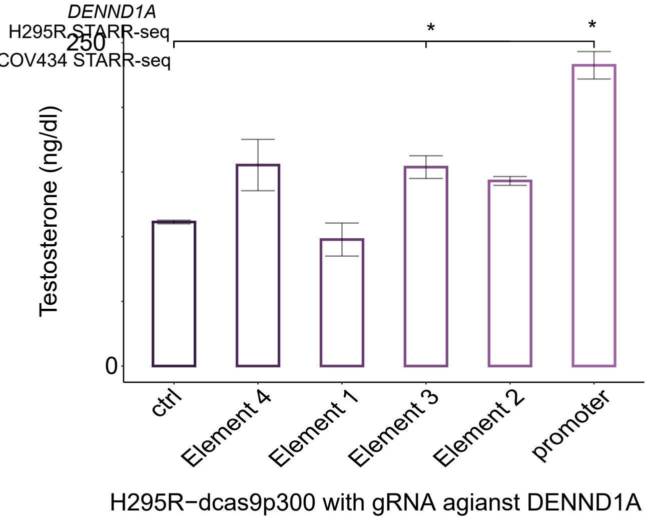

| Primer name | Sequence (5'->3') |
| --- | --- |
| SS_Adaptor_1 | ACACTCTTTCCCTACACGACGCTCTTCCGATCT |
| SS_Adaptor_2 | [Phos]GATCGGAAGAGCACACGTCTGAACTCCAGTCAC |
| TS2SS-F | TAGAGCATGCACCGGACACACTCTTTCCCTACACGACGCTCTTCCGATCT |
| TS2SS-R | GGCCGAATTCGTCGAGTGACTGGAGTTCAGACGTGTGCTCTTCCGATCT |
| 208-F | AATGATACGGCGACCACCGAGATCTACACTCTTTCCCTACACGACGCTCTTCCGATCT |
| Index7 | CAAGCAGAAGACGGCATACGAGAT**NNNNNN**GTGACTGGAGTTCAGACGTGTGCTCTTCCGATC |
| SSRT-UMI | CAAGCAGAAGACGGCATACGAGAT**NNNNNNNN**GTGACTGGAGTTCAGACGTGTGCTCTTCCG*A*T*C |
| Index-PCR | CAAGCAGAAGACGGCATACGA*G*A*T |
| PostSS-Index-5 | AATGATACGGCGACCACCGAGATCTACAC**NNNNNNNN**ACACTCTTTCCCTACAC*G*A*C |
| ssds-F | TAACTTGAAAGTATTTCGATTTCTTGGCTTTATATATCTTGTGGAAAGGACGAAACACCG |
| ssds-R | GTTGATAACGGACTAGCCTTATTTAAACTTGCTATGCTGTTTCCAGCATAGCTCTTAAAC |
| ATAC-universal | AATGATACGGCGACCACCGAGATCTACACTCGTCGGCAGCGTCAGATGTG |
| ATAC-barcode | CAAGCAGAAGACGGCATACGAGAT**NNNNNNNN**GTCTCGTGGGCTCGGAGATGT |
